# Vacidobactin A: An anti- *Pseudomonas aeruginosa* siderophore

**DOI:** 10.64898/2026.02.28.708764

**Authors:** Manpreet Kaur, Derek C. K. Chan, Hodan Wardere, Wenliang Wang, Allison. K. Guitor, Kalinka K. Koteva, Lori L. Burrows, Gerard D Wright

## Abstract

Multidrug-resistant (MDR) *Pseudomonas aeruginosa* poses a significant clinical challenge due to its poorly permeable outer membrane, efflux systems, biofilm formation, and rapid acquisition of resistance genes. The lack of new treatments for *P. aeruginosa* infections underscores the necessity for innovative therapeutic solutions. Iron uptake is essential for bacterial survival, making it a promising target for the development of new antimicrobials. Iron-chelating siderophores are vital for bacterial iron acquisition, important agents for direct antimicrobial action, adjuvants to enhance the effectiveness of currently available antibiotics, and components of prodrugs that facilitate the transport of covalently linked antibiotics into the cell. Here, we report the anti-pseudomonal activity of vacidobactin A, a siderophore produced by the soil bacterium *Variovorax paradoxus,* identified through a screen of natural product extracts targeting a clinical MDR strain of *P. aeruginosa*. Vacidobactin A inhibits *P. aeruginosa* growth by limiting iron availability, particularly in strains that do not produce pyoverdine, their native siderophore. Expression of a TonB-dependent transporter sourced from the vacidobactin producer in a *P. aeruginosa* pyoverdine and pyochelin-null mutant restored its ability to acquire iron and grow in the presence of vacidobactin. Additionally, vacidobactin A synergized with thiostrepton, which hijacks pyoverdine receptors to enter the cell and inhibit protein synthesis. This study supports the therapeutic potential of targeting *P. aeruginosa* iron acquisition pathways and leveraging siderophores as adjuvants to enhance the efficacy of existing antimicrobials. These findings, along with recent advancements in siderophore-based research and combination therapies, offer innovative strategies to combat antibiotic-resistant infections.

**IMPORTANCE:** Multidrug-resistant *Pseudomonas aeruginosa* is a critical priority pathogen for which new therapeutic strategies are urgently needed. Iron acquisition is essential for *P. aeruginosa* survival and virulence, yet remains underexploited as a drug target. Here, we demonstrate that vacidobactin A, a siderophore produced by *Variovorax paradoxus*, suppresses *P. aeruginosa* growth by limiting iron availability, particularly in strains deficient in pyoverdine production. We further show that vacidobactin A enhances the activity of thiostrepton, an antibiotic that exploits siderophore uptake pathways. These findings highlight iron competition as a source of anti-pseudomonal agents and support the development of siderophore-based therapeutics and adjuvant strategies. Targeting iron acquisition networks offers a mechanistically distinct approach to combat antibiotic resistance and expands the repertoire of vulnerabilities that can be leveraged against this highly drug-resistant pathogen.

## INTRODUCTION

Antimicrobial resistance (AMR) is one of the most urgent global health threats, with over one million deaths linked to drug-resistant bacteria in 2021 alone (1). The death toll could exceed 39 million worldwide by 2050 if no action is taken (2), highlighting the need for new therapies. Among the most concerning microbes are Gram-negative bacteria, and among them, *Pseudomonas aeruginosa* is a notoriously challenging opportunistic pathogen. *P. aeruginosa*’s intrinsic antibiotic resistance mechanisms, including a poorly permeable outer membrane, numerous broad-specificity efflux pumps, antibiotic-degrading enzymes, the ability to acquire resistance genes through horizontal gene transfer, and its capacity to form drug-insensitive biofilms, collude to limit effective antibiotic treatment (3, 4).

Despite the increasing prevalence of drug-resistant *P. aeruginosa,* the antimicrobial pipeline targeting this pathogen remains critically underdeveloped. Few of the antimicrobial agents in clinical or preclinical development described in a recent World Health Organization report show anti-pseudomonal activity (5), and murepavadin, a once-promising *Pseudomonas-*specific antibiotic, was discontinued in Phase III clinical trials due to nephrotoxicity (6). Consequently, there is a need to discover new anti-pseudomonal strategies.

Microbially produced natural products offer a valuable source of antimicrobial chemical matter. Indeed, most antibiotics in clinical use are microbial natural products or their semi-synthetic derivatives. Environmental microbes, in particular, are well-known prolific producers of natural products. We have assembled a collection of thousands of environmental microbes and shown that this is a rich source of antibiotics and other bioactive compounds (7). Since *P. aeruginosa* is frequently found in a variety of aquatic and soil habitats, we reasoned that environmental microbes would be a good source of anti-pseudomonal natural products. A screen of a portion of our collection against a clinical strain of multi-drug resistant (MDR) *P. aeruginosa* identified a vacidobactin siderophore. We show that vacidobactin acts as a xenosiderophore that disrupts iron homeostasis in *P. aeruginosa*. In strains that lack pyoverdine production, this disruption leads directly to iron starvation and growth inhibition. Importantly, in pyoverdine-producing strains, vacidobactin does not simply deprive cells of iron but instead triggers an iron starvation response that increases expression of a TonB-dependent siderophore transporter. This transporter is hijacked by the thiopeptide antibiotic thiostrepton, and its increased expression enhances thiostrepton uptake, resulting in strong synergistic antibacterial activity. Through this mechanism, the vacidobactin–thiostrepton combination is broadly effective against diverse *P. aeruginosa* strains.

## RESULTS

### Identification of vacidobactin A from an environmental bacterium

We screened a portion of our collection of natural product-producing microbes (7), focusing on strains derived from a Hamilton, Ontario soil sample that was the source of a previously identified novel lasso peptide antibiotic, lariocidin (8), against *P. aeruginosa* C0292, an MDR strain isolated from sputum. Of >300 methanolic extracts of environmental bacteria isolated from this source, one, MST127, was active against *P. aeruginosa* C0292 (**Supplementary Table S1)**. MST127 was identified as a strain of the betaproteobacterium *Variovorax paradoxus* based on 16S rRNA and whole-genome sequencing analysis. Bioactivity-guided purification of the fermentation broth yielded the active compound with an *m/z* of 752.3320 [M+H] ^+^ determined by high-resolution mass spectrometry (HRMS), consistent with the molecular formula C_30_H_49_N_7_O_15_ (calculated monoisotopic mass: 752.3309) **(Figure 1a)**.

**Figure 1.**
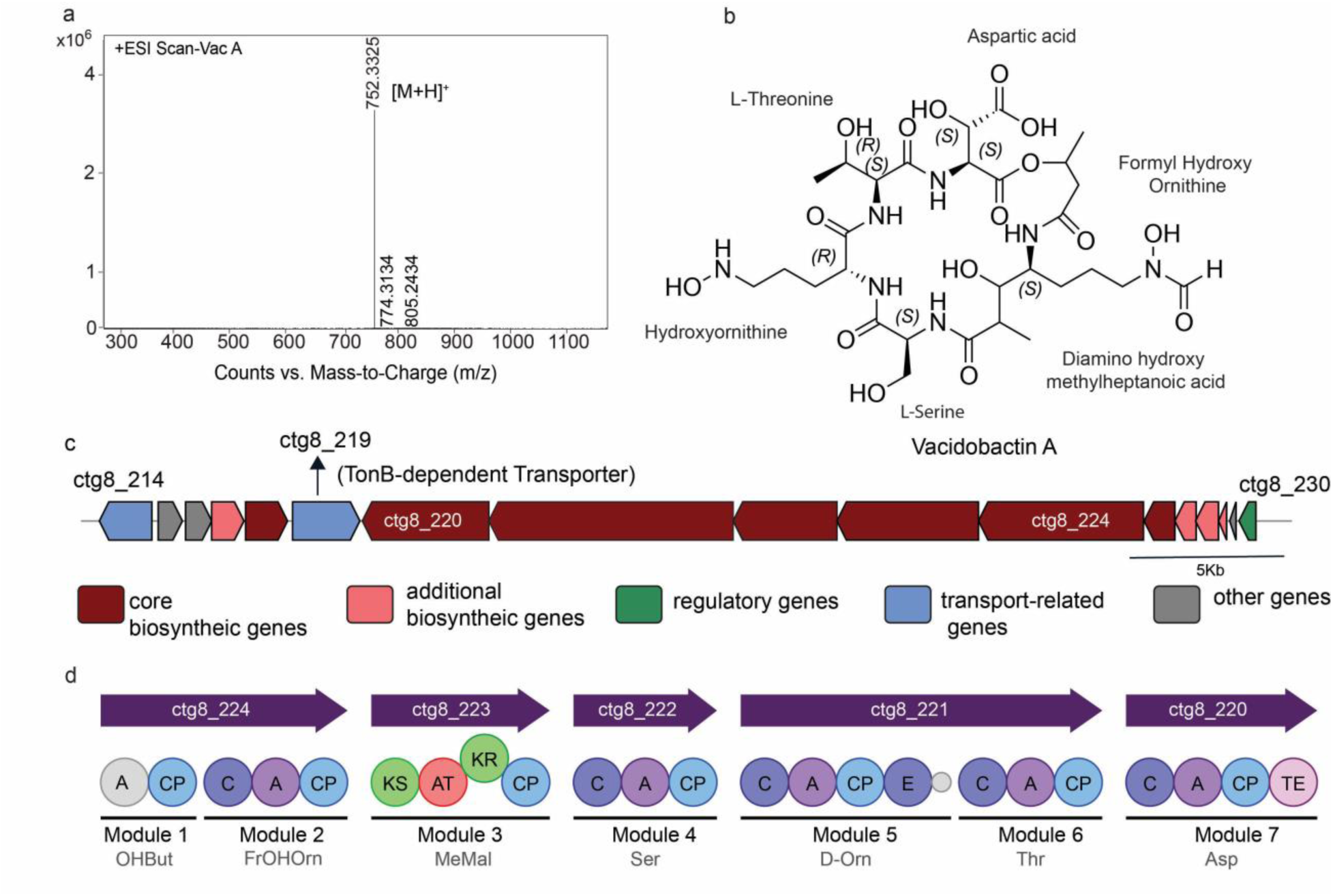
Characterisation of VacA. (a) High-resolution electrospray ionization mass spectrum (HR-ESI-MS) of purified vacidobactin A, confirming the expected molecular mass. (b) Chemical structure of vacidobactin A (Vac-A), annotated to reflect the corresponding amino acid residues. (c) Biosynthetic gene cluster of vacidobactin identified in the whole genome sequence of *Variovorax paradoxus*, as predicted by antiSMASH. Genes are color-coded by function. (d) Domain architecture and predicted amino acid specificity of nonribosomal peptide synthetase (NRPS) modules encoded within the vacidobactin BGC.

The structure of the active compound was determined using one- and two-dimensional nuclear magnetic resonance (NMR) spectroscopy coupled with high-resolution electrospray ionization tandem mass spectrometry (HR-ESI-MS/MS). The compound is a cyclic depsipeptide consisting of five amino acids and 3-hydroxybutyric acid **(Figure 1b).** The latter is linked via an amide bond to an unusual N-terminal 4-amino-3-hydroxy-7-(*N*-hydroxyformamido)-2-methylheptanoic acid. 3-Hydroxybutyric acid provides the alcohol for intramolecular cyclization via an ester bond with the C-terminal β-hydroxyaspartate (β-OHAsp). The presence of the rare 4-amino-3-hydroxy-7-(*N*-hydroxyformamido)-2-methylheptanoic acid residue in the N-terminus suggested similarities to variochelin (9), imaqobactin (10), variobactin (11), and vacidobactin (11) siderophores from *Variovorax* sp. that incorporate modified 4,7-diamino-3-hydroxy-2-methylpetanoic acid elements. Our structural analysis identified the active compound as a member of the vacidobactin family, a group of ß-hydroxyaspartate-acylated siderophores (11). The β-OHAsp and the N-hydroxyornithine provide the key chelating groups necessary for binding Fe (III) ions (12).

The production of vacidobactin A and vacidobactin B has been previously reported from *Variovorax paradoxus* S110 (9). However, their inhibitory activity against *P. aeruginosa* has not been documented. In our initial analyses, the siderophore isolated from the MST127 strain appeared to differ from the published structure of vacidobactin A in the relative positions of Ser and Thr residues, leading us to tentatively designate it as a new variant. To resolve this discrepancy, we obtained the *V. paradoxus* DSM 30334 strain from the original authors and re-isolated vacidobactin A. Detailed NMR and MS analyses demonstrated that the Ser and Thr positions in vacidobactin A from DSM 30334 are identical to those in the compound produced by MST127. This indicates that the previously reported structure of vacidobactin A was misassigned, and that the siderophore we identified in MST127 is, in fact, vacidobactin A with a corrected structure **(Supplementary Figure S1)**. A direct comparison of the MS and NMR spectra from the MST127-derived compound and DSM 30334-derived vacidobactin A is provided in **Supplementary Figure S2–S6.** Additional details of structural assignments, including full NMR spectral datasets, MS, MS/MS analysis, are provided in **Supplementary Figures S7–S16** and **Supplementary Table S5**. The stereochemistry of the proteinogenic amino acids threonine and serine was experimentally confirmed to be L, whereas ornithine is predicted to be in the D-configuration based on the presence of an epimerase domain in its biosynthetic gene cluster **(Supplementary Figure S17).**

AntiSMASH(13) analysis of the *V. paradoxus* MST127 genome identified a putative biosynthetic gene cluster (BGC) containing both polyketide synthase (PKS) and non-ribosomal peptide synthetase (NRPS) components responsible for its production **(Figure 1c and d).** Five core NRPS and PKS-encoding genes, *ctg_220* to *ctg_224*, were identified, providing seven modules predicted to incorporate distinct amino acid and polyketide components. The 4-amino-3-hydroxy-7-(*N*-hydroxyformamido)-2-methylheptanoic acid is presumably generated by *in situ* reaction of NRPS and PKS carrier protein-tethered *N*-formyl-hydroxyornithine and methylmalonic acid components(9). Comparative genomic analysis of the vacidobactin BGCs from MST127 and *V. paradoxus* S110 (DSM 30334) **(Supplementary Table S6)** revealed conserved enzymatic architectures, including seven PKS/NRPS modules, consistent with the production of closely related siderophore scaffolds.

### Antimicrobial activity profile of VacA against selected bacterial strains

VacA exhibited antimicrobial activity against a subset of *P. aeruginosa* strains from our in-house collection of clinical isolates (strains C0063, C0307, C0222, C0157, C0166, C0292, C0090, and C0060) with minimum inhibitory concentrations (MICs) of 16–32 µg/mL in cation-adjusted Mueller-Hinton broth (**Figure 2a**). In contrast, other strains of *P. aeruginosa* (including standard strains such as PAO1 and PA14)*, Acinetobacter baumannii*, and *Klebsiella pneumoniae* were insensitive to VacA, with MICs >128 µg/mL. Time-kill kinetics studies for VacA-sensitive *P. aeruginosa* C0060 showed bacteriostatic activity **(Figure 2b)**, consistent with its siderophore properties, whose antimicrobial activity is governed by iron availability(14).

**Figure 2.**
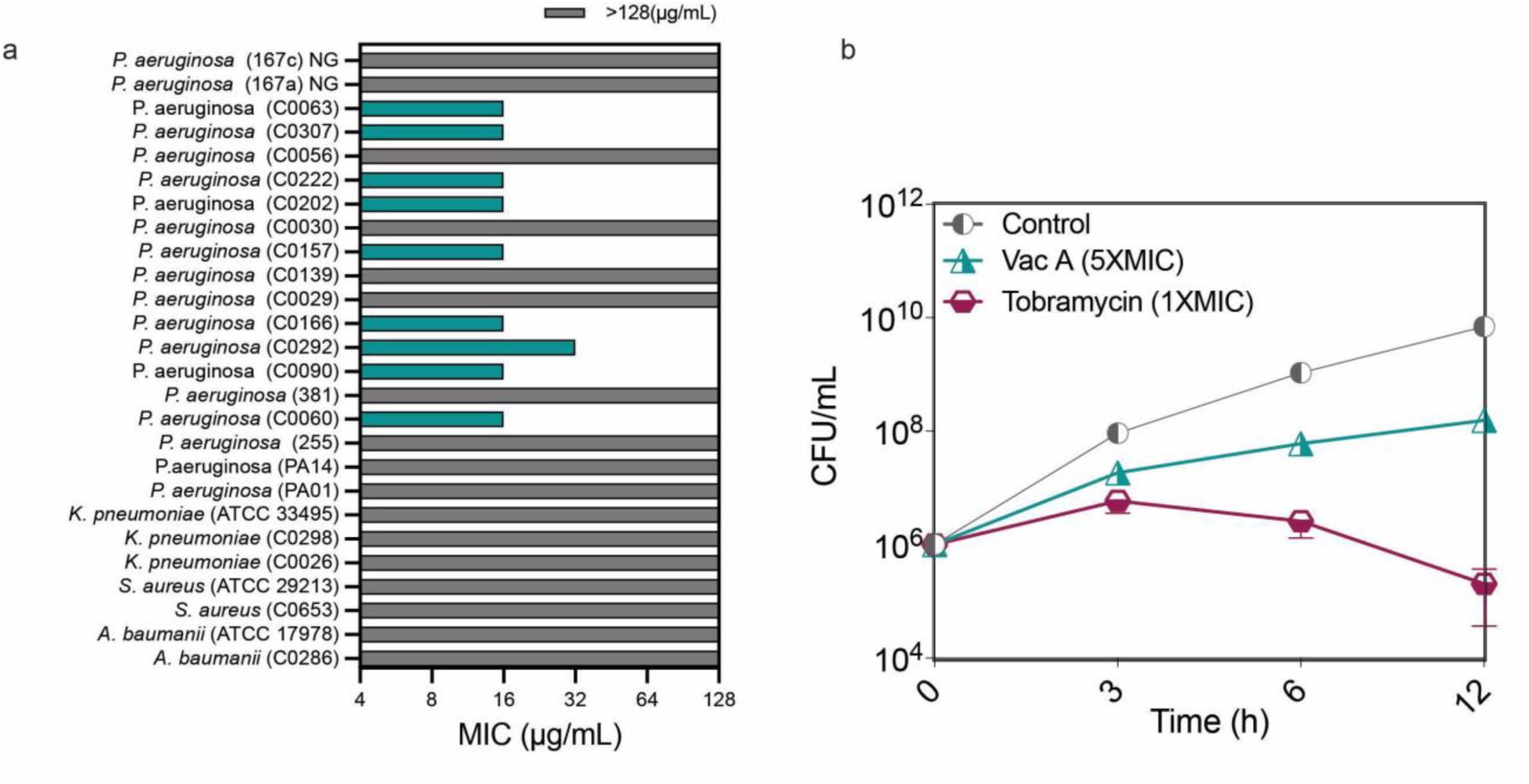
Antimicrobial activity of VacA. (A) Minimum inhibitory concentrations (MICs) of VacA against various bacterial strains, including clinical *Pseudomonas aeruginosa* isolates, determined in cation-adjusted Mueller-Hinton broth (CAMHB). (B) Time-kill kinetics of VacA against *P. aeruginosa* C0060. Bacterial cultures were treated with VacA, and viable counts were determined at the indicated time points. Data represent means ± standard deviation from three independent biological replicates, each performed in duplicate.

### VacA inhibits *P. aeruginosa* via iron starvation in pyoverdine-deficient strains

The role of iron availability in VacA anti-pseudomonal activity was investigated by determining MICs in the presence and absence of supplemental iron (FeCl₃). In iron-limited M9 minimal medium (∼0.6 µM Fe³⁺), the MIC for *P. aeruginosa* C0060 was 8 µg/mL. However, the MIC increased to 16 µg/mL and 32 µg/mL with the addition of 10 µM and 20 µM FeCl₃, respectively, and >128 µg/mL with the addition of 30 µM and 40 µM FeCl₃ **(Figure 3a).** An increase in the MIC of VacA after adding iron to the media suggested that VacA was interfering with iron uptake or metabolism, thereby inhibiting the growth of sensitive strains of *P. aeruginosa*. To investigate this selective sensitivity further, we compared the MICs of VacA-insensitive *P. aeruginosa* PA14 strains with those of mutants from a PA14 transposon library (15). Transposon insertions in *pvdA,* essential for producing the intrinsic siderophore pyoverdine, and *fpvA,* which is the primary pyoverdine receptor, conferred susceptibility to VacA (**Figure 3b)**. Supplementation with pyoverdine alone **(Figure 3c)** or pyoverdine and VacA in combination restored growth, suggesting that pyoverdine competes with VacA for iron. To demonstrate that pyoverdine has superior iron affinity compared to VacA, we performed an assay in which pyoverdine’s intrinsic fluorescence is quenched when binding to Fe^3+^ (16); notably, VacA does not fluoresce (**Figure 3d)**. Pyoverdine supplemented with iron (FeCl_3_, 1 µg/ml) in the presence or absence of VacA showed no significant difference in fluorescence, suggesting it has a higher affinity for iron than VacA (**Figure 3e)**. These results were consistent with the hypothesis that VacA was restricting growth through iron starvation, but only in pyoverdine non-producer strains. We conducted another experiment to evaluate pyoverdine production. A panel of strains isolated from various clinical samples was grown under iron-limited conditions, and fluorescence was measured relative to a positive control (PA14) and a negative control (Δ*pvdA*). As shown in **Supplementary Figure S18**, most isolates produced substantial pyoverdine, while a subset showed minimal or no fluorescence, indicating impaired or absent pyoverdine biosynthesis. Notably, these strains were sensitive to VacA as indicated in **Figure 2a**.

**Figure 3.**
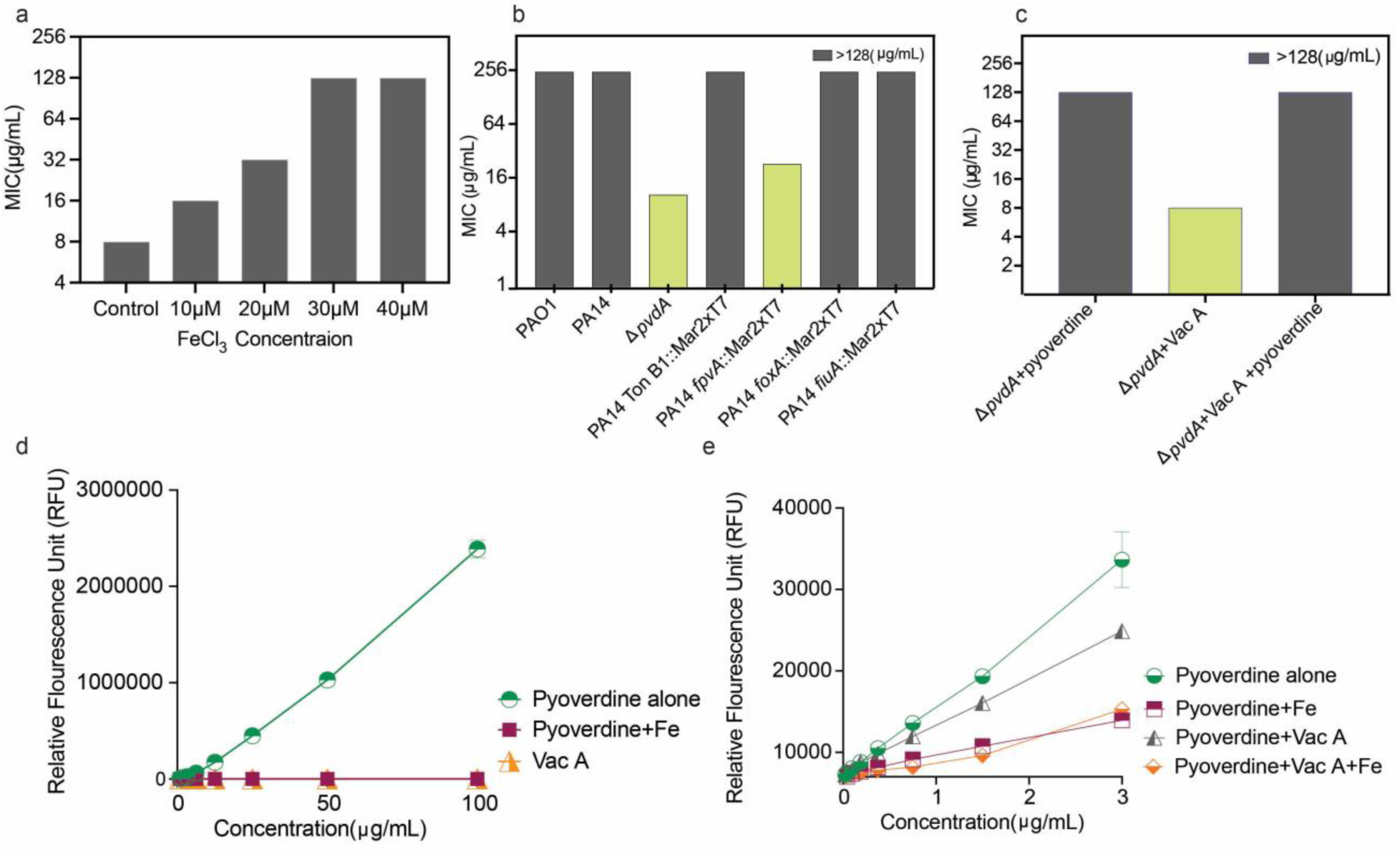
Interaction of VacA with *P. aeruginosa* deletion mutant strains and pyoverdine. (a) Minimum inhibitory concentration (MIC) of VacA against *P.aeruginosa* strain C0060 in M9 minimal medium (baseline iron concentration (0.6 µM)) supplemented with increasing concentrations of FeCl₃, compared to control conditions. (b) MIC of VacA against *P. aeruginosa* PA14 mutant strains defective in iron uptake pathways. (c) Effect of exogenous pyoverdine addition on the MIC of vacidobactin A in the Δ*pvdA* mutant. (d) Schematic representation of pyoverdine fluorescence when unbound to Fe. (e) Fluorescence measurements show that pyoverdine maintains similar fluorescence intensity in the presence or absence of VacA when complexed with Fe, suggesting pyoverdine has a higher iron-binding affinity than VacA. Data represent means ± standard deviation from three independent biological replicates, each performed in duplicate.

### Expression of a TonB-dependent transporter of *V. paradoxus* in *P. aeruginosa* PA14 Δ*pvdA* Δ*pchA* enables VacA- dependent iron uptake

The VacA BGC of *V. paradoxus* MST 127 encodes a predicted TonB-dependent transporter that we hypothesized is responsible for the uptake of the VacA-Fe^3+^ complex **(Figure 1c)**. The TonB-dependent transporter from *V. paradoxus* was expressed in the *P. aeruginosa* PA14 Δ*pvdA* Δ*pchA* mutant to explore whether VacA can be used as an iron source in the absence of native siderophores. This strain lacks the ability to synthesize pyoverdine (due to deletion of *pvdA*) and pyochelin (due to deletion of *pchA*), the two primary siderophores produced by *P. aeruginosa*, rendering it unable to acquire iron via its endogenous siderophore pathways. The ferrichrome transporter, FiuA, from *P. aeruginosa* PA14 most closely resembles the putative *V. paradoxus* VacA transporter and shares 40% identity **(Figure 4a)**. In addition, ferrichrome and vacidobactin share partial structural features characteristic of hydroxamate-type siderophores, providing a rationale for using FiuA as a control transporter in this study. The predicted vacidobactin A transporter could not be functionally expressed in the siderophore-deficient mutant because the N-terminus is important for TonB recognition. Therefore, we generated a chimera that replaced the N-terminus of the *V. paradoxus* TonB-dependent transporter with that of FiuA (PA14_06160) to ensure functionality in *P. aeruginosa* **(Figure 4b).** The control strains *P. aeruginosa* PA14 Δ*pvdA* Δ*pchA* pHERD20T (empty vector) and *P. aeruginosa* PA14 Δ*pvdA* Δ*pchA* pHERD20T-*fiuA* were sensitive to VacA with MIC values of 16 µg/mL, while PA14 Δ*pvdA* Δ*pchA* pHERD20T-N*_fiuA_-*MST 127 was insensitive to VacA with a MIC >128µg/mL**(Figure 4c).**

**Figure 4.**
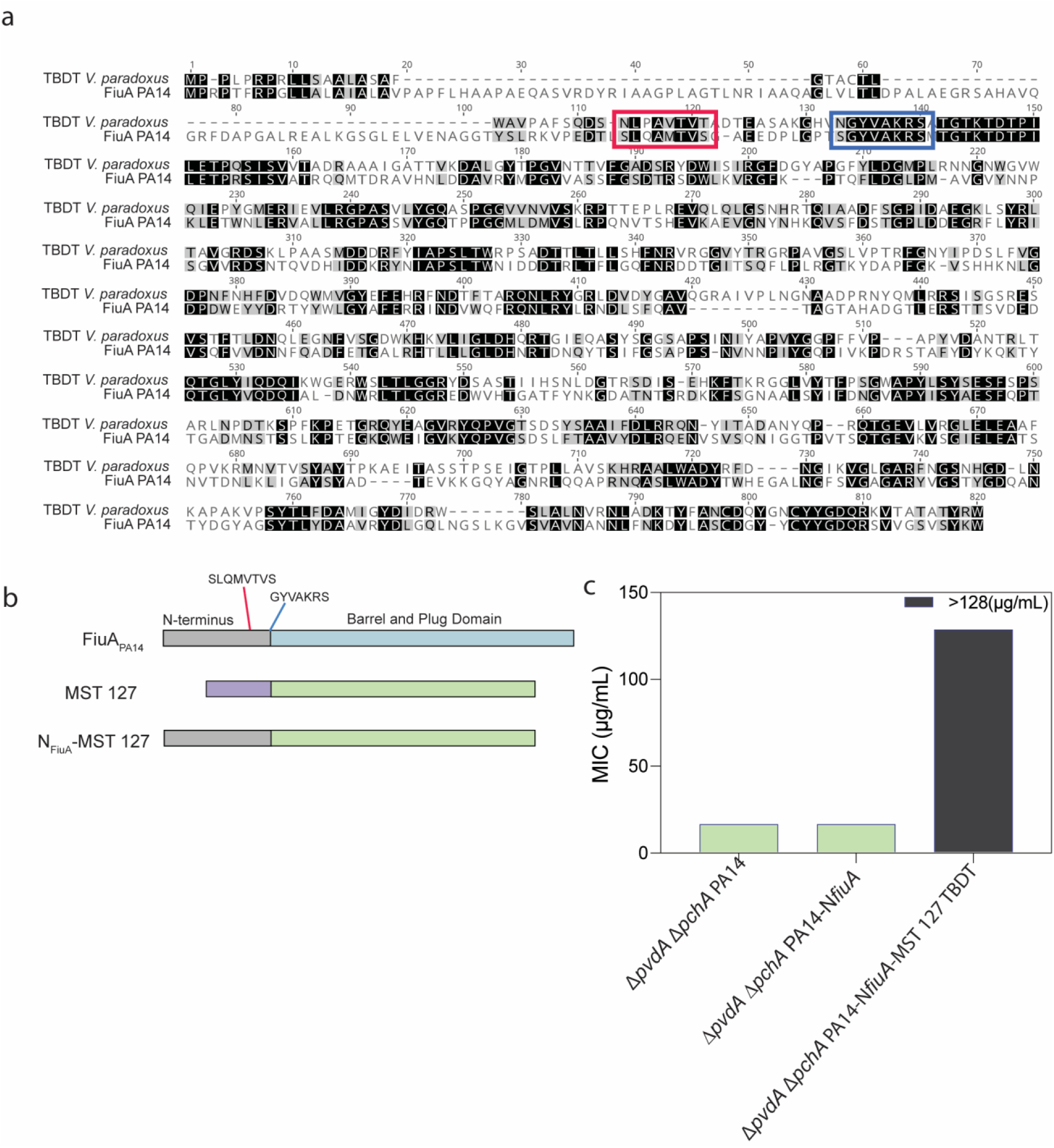
Genetic and functional characterization of VacA biosynthesis and TonB-dependent transport. (a) The sequence alignment of the FiuA TonB-dependent transporter from *P. aeruginosa* PA14 (PA14_06160) and the TonB-dependent transporter from *Variovorax paradoxus* shows 40% identity. The TonB box is highlighted in red, and the overlap region is highlighted in blue. (b) Graphical representation of the fusion of the two transporter sequences, with the TonB box of PA14 FiuA, indicated in red and the overlap region in blue. (c) MIC of mutant strains expressing the Δ*pvdA* Δ*pchA* pHERD20T (empty vector), Δ*pvdA* Δ*pchA* pHERD20T-*fiuA* (vector-fiuA), and Δ*pvdA* Δ*pchA* pHERD20T-N*_fiuA_-*MST 127 TBDT (chimeric transporter N-FiuA-MST 127) the producer strain demonstrates the receptor’s functional role in VacA activity.

These results are consistent with the chimeric transporter facilitating VacA-dependent iron uptake in a manner that supports growth under the conditions tested.

### VacA potentiates the activity of thiostrepton against *Pseudomonas* strains

In our previous work, we showed that iron starvation induces production of the pyoverdine receptors FpvA or FpvB, which unexpectedly facilitated the uptake of large, otherwise impermeant antibiotics such as thiostrepton (17). To evaluate the potential of VacA as an antibiotic adjuvant, we performed synergy assays to measure the combined effect of thiostrepton and VacA against *P. aeruginosa*, including standard strains (PAO1, PA14) and clinical isolates, including pyoverdine producers (C0062, C0030) and non-producers (C0060, C0090). Synergistic interactions were observed for all tested strains, with pronounced synergy in pyoverdine-producing isolates and reduced synergy in non-producers, likely reflecting their individual MICs **(Figure 5a-f).**

**Figure 5.**
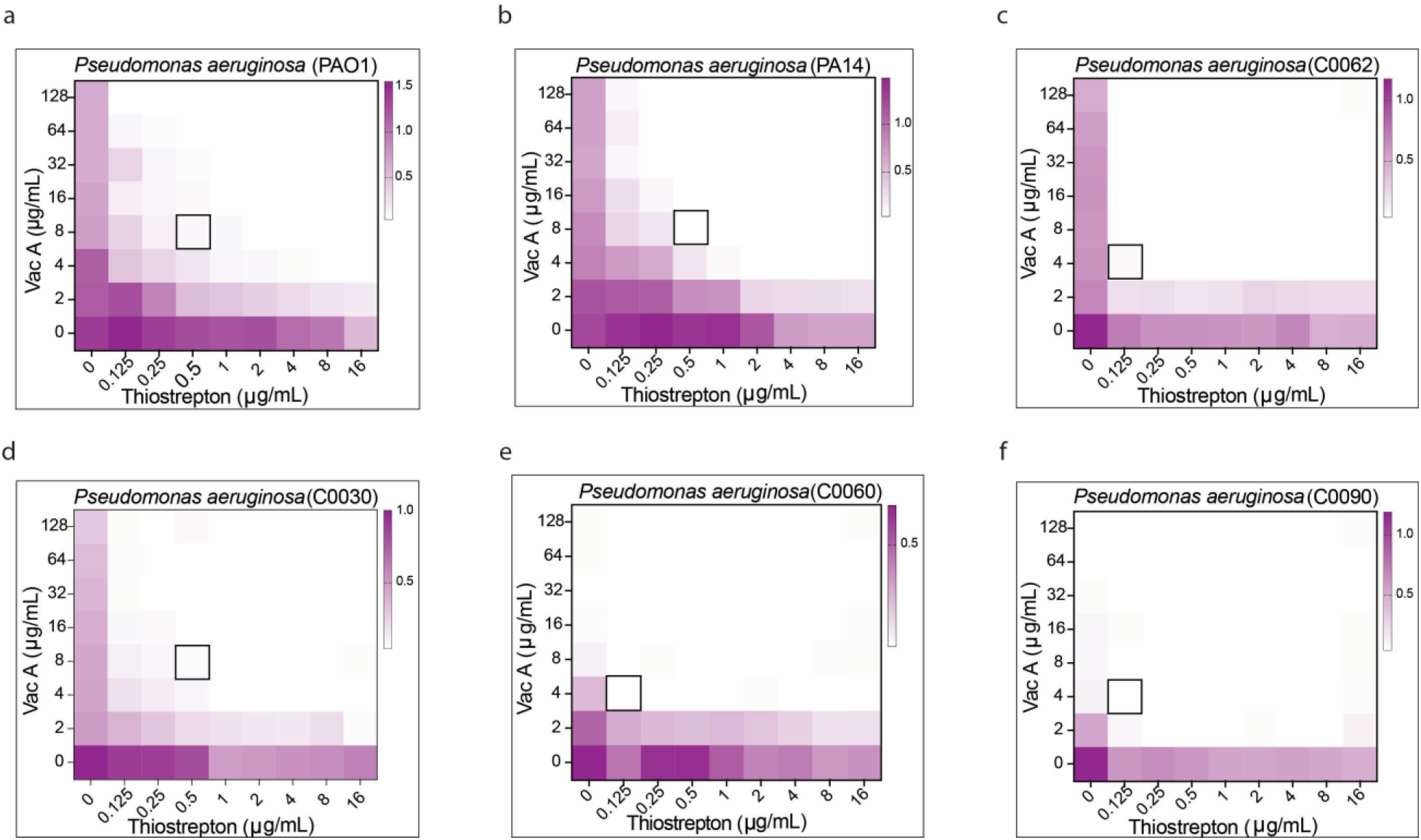
Synergistic antibacterial activity of VacA and thiostrepton against *P. aeruginosa* strains, including clinical isolates. (a–f) Synergy assay heatmaps illustrating the combined effects of thiostrepton and VacA against *P. aeruginosa* strains, including laboratory strains PAO1 and PA14, and clinical isolates C0062 and C0030 (pyoverdine producers), and C0060 and C0090 (non-pyoverdine producers). Concentrations of thiostrepton (x-axis) and VacA (y-axis) are indicated for each panel. Growth inhibition is represented by shading intensity, with lighter colors indicating greater inhibition. Regions of synergistic interaction, where subinhibitory concentrations of both compounds lead to pronounced growth inhibition, are highlighted with black boxes. The data shown represent one of three independent biological replicates, each yielding results comparable to the others.

The combination of VacA (8 µg/mL) and thiostrepton (0.5 µg/mL) significantly suppressed growth of a panel of 21 clinical *P. aeruginosa* isolates. As shown in **Supplementary Figure S19**, the extent of inhibition varied among isolates, but overall, the combination consistently reduced bacterial growth. These findings highlight the efficacy of VacA and thiostrepton as a potential antimicrobial combination, particularly against pyoverdine-producing *P. aeruginosa* strains.

### VacA enhances pyoverdine production and increases receptor availability for thiostrepton uptake

To assess the effect of VacA on pyoverdine production, we treated *P. aeruginosa* PA14 with VacA (8 µg/mL), thiostrepton (0.5 µg/mL), or their combination. Optical density measurements (OD₆₀₀) revealed that PA14 grew well under control conditions and showed minimal growth inhibition with thiostrepton alone. However, VacA treatment significantly reduced growth, with OD₆₀₀ values approximately half of the control **(Figure 6a).**

**Figure 6.**
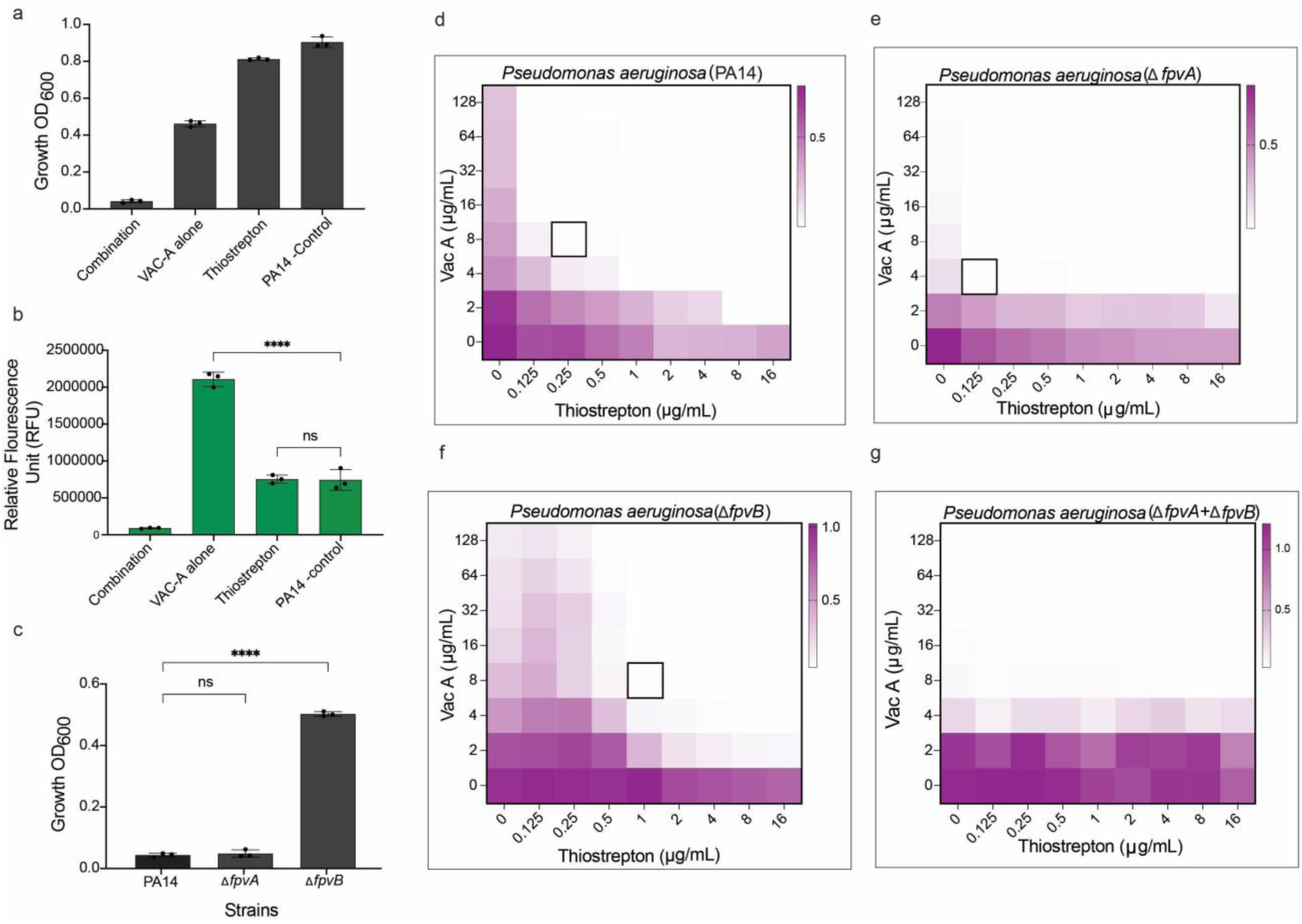
VacA enhances pyoverdine production and promotes thiostrepton uptake via siderophore receptors. (a) Growth (OD₆₀₀) of *P. aeruginosa* PA14 after 18 h treatment with VacA (8 µg/mL), thiostrepton (0.5 µg/mL), or a combination of the two. b) Pyoverdine fluorescence following treatment, showing increased pyoverdine production in response to VacA alone. (C) Growth inhibition of wild-type PA14 and the Δ*fpvA* mutant in combination, demonstrating increased sensitivity to the combination in comparison to the Δ*fpvB* mutant. (d-g) Synergy between VacA and thiostrepton in the deletion mutant strains. Data represent the mean ± SD of at least three independent experiments. P values were determined using Dunnett’s multiple comparisons test; P < 0.0001 for both (b) and (c)

Next, we examined pyoverdine production under the same conditions. VacA alone markedly induced pyoverdine synthesis, whereas thiostrepton had no effect **(Figure 6b).** These results suggest that VacA enhances pyoverdine production, potentially increasing the availability of siderophore receptors needed for thiostrepton uptake. In *P. aeruginosa,* ferripyoverdine binding to the outer membrane receptor FpvA activates the extracytoplasmic function sigma factor PvdS, which coordinately upregulates both pyoverdine biosynthesis genes and *fpvA* expression, thereby establishing a positive feed-forward regulatory loop(18). Increased pyoverdine production would therefore be expected to increase FpvA receptor abundance, potentially enhancing thiostrepton uptake.

To test this hypothesis further, we evaluated the synergy between VacA and thiostrepton in PA14 mutant strains lacking either the pyoverdine receptors FpvA or FpvB. The Δ*fpvB* mutant showed reduced sensitivity to the VacA–thiostrepton combination, indicating that higher concentrations are required for synergy in the absence of FpvB. In contrast, the Δ*fpvA* mutant and Δ*fpvA*-Δ*fpvB* behaved similarly to pyoverdine-deficient strains, displaying sensitivity to VacA alone suggesting FpvA and FpvB are critical for iron acquisition in the presence of VacA. Comparative synergy analyses in these mutant strains are presented in **Figures 6d–g**.

## Discussion

This study demonstrates that VacA, a siderophore produced by environmental *V. paradoxus*, exhibits selective antimicrobial activity against pyoverdine-deficient *P. aeruginosa* strains through iron starvation. Furthermore, VacA shows a potent synergistic effect with the large-scaffold antibiotic thiostrepton across a broad range of *P. aeruginosa* isolates regardless of their pyoverdine production status. VacA was originally isolated and characterized by Johnson et al (11) from *V. paradoxus* S110 strain, however, no antimicrobial activity was reported. This may be due to the absence of VacA activity against standard *P. aeruginosa* strains, such as PAO1 and PA14, which are often used for antibacterial screening. We identified VacA during a targeted screen of a multidrug-resistant *P. aeruginosa* clinical isolate, C0292, which does not produce pyoverdine. This observation highlights the impact of genetic variability among clinical isolates, including collateral sensitivity, in which resistance to one antibiotic may confer hypersensitivity to an unrelated compound (19). These results also emphasize the critical role of indicator strain selection in antimicrobial discovery efforts and support the inclusion of clinically relevant, resistant isolates alongside standard strains in screening pipelines. Our structural re-characterization of VacA corrects the previously misassigned positions of serine and threonine residues in the originally reported structure. This correction, confirmed through comparative analysis with the reference strain DSM 30334, highlights the importance of rigorous structural validation in natural product research.

Iron acquisition is a central determinant of *P. aeruginosa* fitness in both environmental and host-associated settings, including in biofilm formation, where iron is frequently limiting and tightly sequestered (20–22). To survive under iron-limiting conditions, *P. aeruginosa* produces the high-affinity iron-chelating compounds pyoverdine and pyochelin. The selective activity of VacA against pyoverdine-deficient *P. aeruginosa* strains reveals an important vulnerability in this notoriously drug-resistant pathogen. Our findings demonstrate that VacA’s inhibitory activity results from competitive iron chelation, effectively starving bacteria that cannot produce their primary siderophore, pyoverdine. This iron-dependent activity is evidenced by the dose-dependent reversal of growth inhibition upon iron supplementation and the restoration of growth when pyoverdine is provided exogenously. The selective nature of VacA’s activity aligns with the ecological context in which it is produced. *V. paradoxus*, commonly found in soil and aquatic environments where *P. aeruginosa* also thrives, may have evolved VacA production to provide a competitive advantage for iron acquisition in these shared habitats. These results align with previous reports demonstrating the antimicrobial activity of siderophores, such as variochelins, produced by various *Variovorax* species (23). While previous studies did not specifically test variochelins against *P. aeruginosa*, the demonstrated antimicrobial activity against other Gram-negative bacteria aligns with our findings and suggests that *Variovorax*-derived siderophores may represent a broader class of competitive iron-chelating antimicrobials. The structural similarities between variochelins and VacA, particularly the presence of β-hydroxyaspartate, and N-hydroxyornithine chelating groups, likely contribute to their shared ability to disrupt bacterial iron homeostasis.

Siderophores produced by *P. aeruginosa* require specific TonB-dependent transporters to cross the outer membrane and access to the periplasm and cytoplasmic membrane for further transport into the cell (24, 25). Several antimicrobial natural products can act as Trojan horses, exploiting these iron transporters to enter the cell. For instance, the bulky hydrophobic thiopeptides thiocillin and micrococcin hijack the ferrioxamine receptor FoxA (26), while thiostrepton exploits the pyoverdine receptors FpvA and FpvB (27). Iron chelators potentiate these antimicrobial compounds by inducing iron-limiting conditions, thereby increasing the expression of relevant siderophore transporters. This phenomenon underlies the efficacy of VacA as an adjuvant with the antibiotic thiostrepton. Importantly, VacA demonstrated stronger synergistic activity with thiostrepton compared with previous studies that explored other siderophores in combinations (17, 26, 28). Moreover, the synergistic effect was retained in nutrient-rich conditions (Mueller-Hinton Broth), whereas other studies utilized iron-limiting conditions for the combination. Our data robustly support this established model, which uses thiostrepton or other antibiotics that utilize this uptake mechanism in combination with iron chelators. Targeting iron uptake mechanisms via VacA, either alone or in combination with thiostrepton, appears to be a promising strategy to control *P. aeruginosa* infections and warrants further studies to confirm its potential as an innovative solution to the AMR crisis.

## DATA AVAILABILITY STATEMENT

The complete genome sequence of *Variovorax paradoxus* WAC MST 127 is deposited in NCBI GenBank with BioProject ID SUB15804891

## ACKNOWLEDGMENTS

We thank Dr. Nathan A. Magarvey for providing the *V. paradoxus* DSM 30334 strain used to isolate vacidobactin A. This research was funded by the Canadian Institutes of Health Research (CIHR grants FRN-148463 and PJT183745 to GDW) and by the Natural Sciences and Engineering Research Council of Canada (grant RGPIN-2021-04237 to LLB). LLB holds the Canada Research Chair in Microbe-Surface Interactions (CRC-2021-00103) and DCKC held a CIHR CGS-D award.

## AUTHOR CONTRIBUTIONS STATEMENT

M.K. and G.D.W. conceived the study. M.K. performed the isolation of strains from soil, antimicrobial activity screening, and purification of Vacidobactin A; D.K.C. performed the expression of TonB-dependent transporter *V. paradoxus* in PA14; M.K. and H.W. performed the biochemical and microbiological experiments, including MICs, checkerboard, and synergy assays; W.W conducted the structural analysis and NMR experiments; A.K.G. performed the whole genome sequencing analysis and assembly of the genome. K.K performed the Marfey’s analysis of VacA. M.K. and G.D.W prepared the manuscript with input from other authors. L.L.B and G.D.W. supervised the study and edited the manuscript. All authors approved the manuscript before submission.

## Material and Methods

### Bacterial strains and culture conditions used in this study

The strains, plasmids, and oligonucleotide primers used in this study are listed in **Supplementary Tables S2-S4**. *V. paradoxus* strain MST 127 was grown in ISP2 medium (Yeast extract (Difco) 4.0 g L, Malt extract (Difco) 10.0 g L, Dextrose (Difco) 4.0 g L, pH 5.6) with 1 % glycerol and in a shaking incubator at 30°C for 48h. MIC experiments were performed in cation-adjusted Mueller Hinton Broth (MHB; BD Biosciences), modified M9 medium (6.8 g L, Na_2_HPO_4_, 3 g L, KH_2_PO_4_, 0.5 g L, NaCl, 1 g L’ NH_4_Cl, 0.4% glucose, 2 mM MgSO_4_, 0.1 mM CaCl_2_, 0.2% casein amino acids,16.5 μg mL–1 thiamine)^13^ and CAA (casamino acid medium, composition: 5 g L^−1^low-iron CAA (Difco), 1.46 g L^−1^ K_2_HPO_4_ 3H_2_O, and 0.25 g L^−1^ MgSO_4_ 7H_2_O) (29). Lennox Broth (LB; Bioshop) was used to test the intracellular iron level experiments. All strains were incubated at 37°C.

### Screening of soil isolates and activity-guided purification of vacidobactin A

Bacterial strains isolated from soil samples were screened for their antimicrobial activity against the multidrug-resistant clinical strain of *P. aeruginosa* C0292. Tryptic soy broth TSB, half-strength Brain Heart Infusion (BHI) broth, and minimal media supplemented with peptones were selected for soil isolate fermentation. One ml of cell-free broth was collected after 24 h, 48 h, 5 days, and 10 days of incubation, lyophilized overnight, and resuspended in 50% DMSO (50-70 µl). For antimicrobial activity screening, *P. aeruginosa* was inoculated on Mueller-Hinton agar (MHA) plates, 5 µl of the DMSO extract was applied, and plates were incubated at 37°C overnight. The extract from *V. paradoxus* MST 127 grown for 48 h in minimal media supplemented with peptones showed consistent activity against *P. aeruginosa*. ISP2 medium with 1% glycerol was selected for further purification as it yielded the highest antimicrobial activity. A 50 ml seed culture was used to inoculate 3 liters of medium. After two days of incubation at 30°C, the supernatant was separated from the cells by centrifugation at 3400 x g for 20 min. The cell-free supernatant was added to 8% (w/v) of activated Diaion HP-20 (Sigma) resin and mixed for 2-3h. The resin was filtered and washed with distilled water. The compounds bound to the resin were eluted with 100% methanol after 2h of mixing, and the solvent was removed using a rotary evaporator. The extract was reconstituted in Milli-Q water, loaded onto a Sephadex LH-20 column, and eluted with a water-methanol gradient from 0% to 50% methanol. The antibiotic activity of these fractions was assessed using the well-diffusion method. The active fractions were pooled and loaded on a reverse-phase CombiFlash ISCO column (RediSep Gold Rf C18, Teledyne). The compounds were eluted with acetonitrile (ACN) and water using a 0% to 95% ACN gradient. The active fractions were dried and then applied to an HPLC C28 Analytical column (Sunniest RP-AquA C28 4.6X 100 mm, 5µm) using water and eluted with a gradient of ACN. Both solvents were supplemented with 0.1% trifluoroacetic acid. The fractions were collected and assessed for antimicrobial activity. The active peak eluted at approximately 7% ACN. The pure compound, vacidobactin A, was lyophilized to generate a white powder. A similar purification procedure was followed to isolate VacA from the DSM 30034 strain.

### Structural characterization of VacA

The mass and MS/MS data of VacA were analyzed using HR-ESI-MS recorded with an Infinity II LC System (Agilent Technologies) coupled with a qTOF 6550 mass detector in positive ion mode. For NMR spectroscopy, 8mg of pure VacA from MST 127 and 5mg from DSM 30034 were dissolved in deuterated DMSO, and 1D and 2D NMR spectra were acquired on a Bruker AVIII 700 MHz instrument equipped with a cryoprobe.

### Determination of the stereochemistry of VacA amino acids by Marfey analysis

To a solution of VacA (0.6mg) was added 6N HCl (1ml). Peptide bond hydrolysis was carried out for 20h at 90°C with stirring in a closed glass vessel. The sample was then diluted with water and freeze-dried. To the hydrolyzed dry peptide was added water (10µl), followed by 1M sodium bicarbonate (10µl) and (1-fluoro-2,4-dinitrophenyl-5-L-alanine amide) (Marfey’s reagent) (50µl of 1% solution in acetone). The reaction was carried out for 1h at 40 °C and terminated by the addition of 1M HCl (10µl) and methanol (420µl), followed by LC-MS analysis(30).

### Marfey analysis of reference serine and threonine amino acids

To 10µl of 10mg/ml solution of the corresponding amino acid L or DL was added 10µl of 1M sodium bicarbonate (10µl) and Marfey’s reagent (50µl of 1% solution in acetone). The reaction was carried out as described above. High-resolution electrospray ionization mass spectra were acquired using an Agilent 1290 UPLC separation module and a qTOF 6550 mass detector in positive ion mode. For LC separation an Agilent Eclipse XDB-C8 column (2.1 x 150 mm; 3.5 µm and the following method were used: from 0 to 0.5min 90% A (0.1 v/v formic acid in water), 10% B (acetonitrile), from 0.5min to 1-up to 20%B followed by a linear gradient from 1min to 20min to 30% B, then from 20 to 25 min sharp increase to 90%B at a flow rate of 0.2 mL/min.

### Biosynthetic gene cluster identification

The genome sequence of *V. paradoxus* MST 127 was obtained by the hybrid assembly of Illumina MiSeq reads and Nanopore data. The VacA biosynthetic gene cluster was identified using antiSMASH 5.0.(13).

### Minimal inhibitory concentration (MIC) assay

MICs were determined using the CLSI broth microdilution method (31) in a final volume of 100 µl in 96-well U-bottom plates. Cation-adjusted Mueller Hinton Broth (CA-MHB) and M9 minimal media were used for other MIC experiments as indicated. Bacteria were grown overnight, and an inoculum of 5 x 10^5^ CFU/ml (final concentration) was prepared. MICs were also determined in the presence and absence of iron salts. M9 medium was used to prepare the inoculum, and four different concentrations of FeSO_4_ and FeCl_3_ (10, 20 M, 30, and 40 µM) were used.

### Time-dependent killing and minimum bactericidal concentration (MBC) determination

Clinical isolate *P. aeruginosa* C0060 was grown overnight in CA-MHB, and the OD_600 was_ adjusted to 0.1. The cells were diluted 100-fold in fresh media containing 1×MIC and 5×MIC of vacidobactin A or 1×MIC of tobramycin and were incubated at 37°C. The cells without supplemented antibiotics served as the positive control. Ten µl of treated cells were spotted onto CA-MHB agar plates at four different time points (0 h, 3 h, 6 h, and 24 h), and CFUs were counted. The experiment was conducted in duplicate and repeated three times on separate days.

### Generation of *P. aeruginosa* mutants

A *P. aeruginosa* PA14 Δ*pvdA* Δ*pchA* double mutant was generated by allelic exchange. Primers flanking the upstream (∼1000bp) and downstream (∼750bp) regions of *pvdA* and *pchA* were PCR-amplified from PA14 genomic DNA (Promega Wizard Genomic DNA Purification Kit) with the primers described in **Supplementary Table S4** and gel-extracted with GeneJet Gel Extraction Kit (ThermoFisher).

To make the Δ*pvdA* deletion construct, the upstream fragment was digested with EcoRI and BamHI (FastDigest, ThermoFisher) and then ligated into pEX18Gm digested with the same enzymes (T4 DNA ligase, ThermoFisher). The ligation mixtures were transformed into competent *E. coli* DH5α by heat shock with a recovery period of three hours in lysogeny broth (LB, BioShop). Cells were plated on LB 1.5% agar containing 15µg/mL gentamicin supplemented with X-gal for blue-white screening. The plates were incubated at 37°C overnight. White colonies were selected and grown in LB + 15µg/mL gentamicin. Cells were spun down, and the plasmids were isolated using GeneJet Plasmid Miniprep Kit (ThermoFisher). Plasmids were checked by digestion with EcoRI and BamHI. The plasmid was then digested with BamHI and HindIII and ligated with the digested downstream fragment, and the same checks were repeated.

To generate the Δ*pchA* deletion construct, the upstream and downstream fragments were joined by overlap extension PCR and digested with EcoRI and HindIII. pEX18Gm was digested with the same enzymes, and the fragments were ligated overnight with T4 DNA ligase as outlined above.

Plasmids were transformed into competent *E. coli* SM10 by heat shock with a recovery period of 3h in LB. The cells were plated on LB 1.5% agar containing 15µg/mL gentamicin and grown overnight at 37°C. One colony was inoculated in LB + 15µg/mL gentamicin overnight. PA14 was also inoculated from a single colony into LB. Both cultures were grown overnight at 37°C. SM10 pEX18Gm-*ΔpvdA* was mated with PA14 by mixing 500µL of each overnight culture. Cells were centrifuged, and the supernatant was removed. Cells were resuspended in 50µL fresh LB, spotted on LB 1.5% agar, and incubated overnight at 37°C. Cells were streaked onto *Pseudomonas* Isolation Agar (PIA, Difco) supplemented with 100µg/mL gentamicin and incubated overnight at 37°C. Single colonies were streaked onto LB + 15% sucrose (BioShop) and incubated overnight at 37°C. To check for colonies with the correct deletion, 16 colonies were patched onto LB + 15% sucrose and LB + 30µg/mL gentamicin and incubated overnight at 37°C. Patches that grew on the sucrose plates but not gentamicin plates were colony PCR checked with primers flanking the *pvdA* gene compared to wild-type controls. PA14 Δ*pvdA* was streaked onto LB + 15% sucrose to isolate single colonies, incubated overnight at 37°C, and verified by colony PCR. A single colony was inoculated into LB broth, and the process was repeated to delete *pchA* and generate a siderophore-null *P. aeruginosa* mutant.

### Growth restoration assay between pyoverdine and VacA

The MIC of VacA was determined in M9 medium for a *P. aeruginosa* Δ*pvdA* mutant (unable to synthesize pyoverdine (32)) in the presence and absence of pyoverdine (10 µM) (Sigma Aldrich). The difference in the MICs of VacA in the presence and absence of pyoverdine was used to measure competition between the two siderophores.

### Comparison of iron affinity between VacA and pyoverdine

This experiment was conducted to compare the affinity of VacA and pyoverdine for iron. Apo-pyoverdine fluoresces with excitation at 407 nm and emission at 460 nm, which is quenched when it binds to Fe^3+^. Pyoverdine or VacA were 2-fold serially diluted from 100 µg/mL to 1.2 µg/mL in black 96-well microplates. FeCl_3_ (100 µg/mL) was added to pyoverdine-containing wells to quench fluorescence. The fluorescence was recorded immediately using a SynergyH1 microplate reader (excitation/emission 407 nm/460 nm). VacA has no intrinsic fluorescence. Next, the effect of vacidobactin on the fluorescence of pyoverdine was determined in the presence or absence of Fe^3+^. VacA and pyoverdine were added in combination or alone at 1-3 µg/ml. FeCl_3_ was kept constant at 1 µg/mL, and the fluorescence was recorded as described (16).

### Measurement of growth and pyoverdine production levels

To determine whether the clinical isolates could produce pyoverdine, we cultured all isolates under iron-limited conditions and assessed pyoverdine production. Initially, the isolates were grown in 150μl of LB medium in 96-well plates under static conditions at a 37 °C incubator for 16–18 hours. The next day, 2μl of the overnight cultures were transferred to 200μl of iron-limited CAA medium in 96-well plates. The CAA medium was supplemented with 25mM HEPES buffer, 20mM NaHCO₃, and 100μg/ml human apo-transferrin, a potent natural iron chelator. The experiment was performed in triplicate, with Δ*pvdA* as the negative control and PA14 as the positive control. After incubating the plates at 37°C incubator in the dark for 18h, growth was measured as optical density at 600 nm (OD_600_), and pyoverdine production was quantified as relative fluorescence units (RFU) using a BioTek Synergy plate reader (excitation at 400nm and emission at 460nm). Relative growth and pyoverdine production for each isolate were calculated by normalizing their OD600 and RFU values to the average OD600 and RFU of the reference strains (14).

### Preparation of overexpression strains

Primers were used to amplify full-length *fiuA* (*PA14_06160*) plus 22 bp upstream of the start codon encompassing its putative translational start site. The sequence encoding the N-terminus was also amplified (417 bp after the start codon) with a 20 bp extension for the overlapping region of the *V. paradoxus* TonB-dependent transporter in the BGC predicted by antiSMASH. Similarly, the overlapping region from the *V. paradoxus* TonB-dependent transporter was amplified from *V. paradoxus* genomic DNA to the stop codon. The N-terminus of FiuA was fused to the barrel and plug domain of the *V. paradoxus* TonB-dependent transporter by overlap extension PCR. The chimeric transporter was expressed from the arabinose-inducible plasmid, pHERD20T (33).The full-length fusion fragment and *fiuA* were digested with EcoRI and HindIII (FastDigest, ThermoFisher) and ligated into pHERD20T, also digested with EcoRI and HindIII.. Ligations and selection protocols were identical except for the blue-white screening method; plates were also supplemented with 100 µL of 20% arabinose in LB. Correct plasmids were electroporated into PA14 Δ*pvdA* Δ*pchA* with a recovery period of 1 h and plated onto LB 1.5% agar + 200 µg/mL carbenicillin. Plates were incubated overnight at 37°C and single colonies were selected for MIC assays.

### Checkerboard synergy assays

The synergy assay for VacA and thiostrepton was conducted using a checkerboard format in 96-well plates. An 8-by-9 checkerboard layout was used, with the concentration of one compound increasing along the y-axis and the concentration of the second compound increasing along the x-axis. The assay was performed in cation-adjusted Mueller-Hinton broth (CaMHB). Each well had a final volume of 100 µL, and serial dilutions of the compounds were prepared to achieve a range of concentrations for each drug. Each well was inoculated with *P. aeruginosa* culture at a final inoculum of 5 × 10^5 CFU/mL. The plates were covered to prevent evaporation and incubated at 37°C for 16-18h hours. Following incubation, the optical density at 600 nm (OD_600_) was measured using a BioTek microplate reader. At least three biological replicates were performed for each combination. Data were analyzed in Excel and plotted against compound concentrations using Prism software (GraphPad).

### Combination synergy assay

To further evaluate the synergistic effect of VacA and thiostrepton, a combination assay was performed using 21 clinical *P. aeruginosa* isolates. Freshly streaked plates were used to inoculate 200 µL of CaMHB broth, followed by incubation for 16 h at 37°C. After overnight incubation, 6 µL of the culture was transferred into 144 µL of fresh CaMHB and incubated for 2-4 h at 37°C to reach mid-log phase. For the combination assay, each well contained 2 µL of the 2-4h subculture of *P. aeruginosa*. For the assay, 2 µL of a combination of VacA (8 µg/mL) and thiostrepton (0.5 µg/mL) and 146 µL of Cation adjusted-MHB media, for a total volume of 150 µL per well. The plates were incubated overnight at 37°C. After 16 h, the OD_600_ was measured to assess bacterial growth inhibition using a plate reader. Each experiment was conducted in triplicate and with three biological repeats to ensure reproducibility

## SUPPLEMENTARY FIGURES

**Figure S1.**
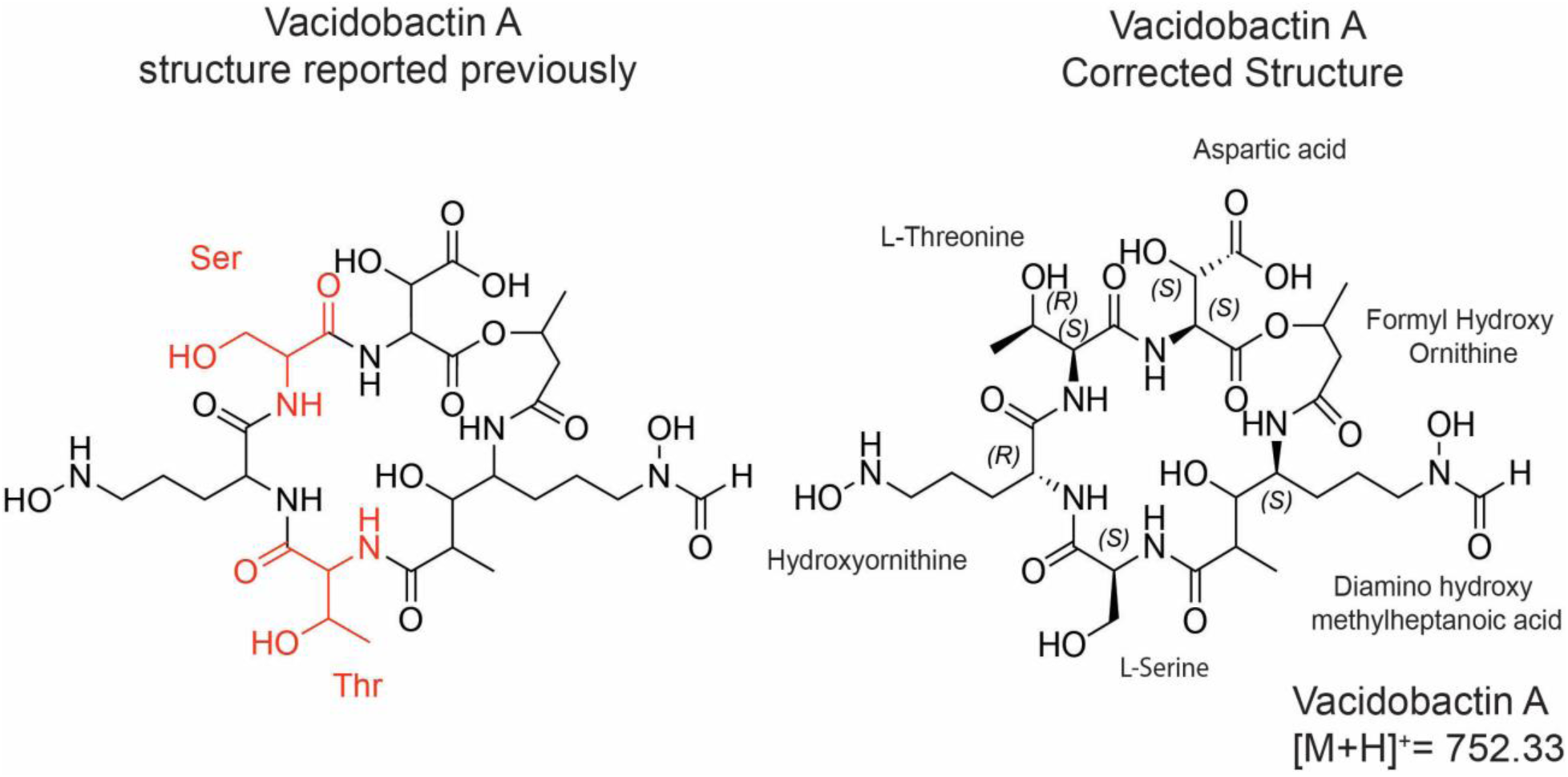
Structural differences between Vacidobactin A (previously reported) and the corrected Vacidobactin. **A.** Key structural variations are highlighted in red in the previously reported Vacidobactin A. In the corrected Vacidobactin A, a peptide bond is formed between β-OHAsp and threonine at position 2. In contrast, the previously reported Vacidobactin A features a serine at position 2, shown in red

**Figure S2.**
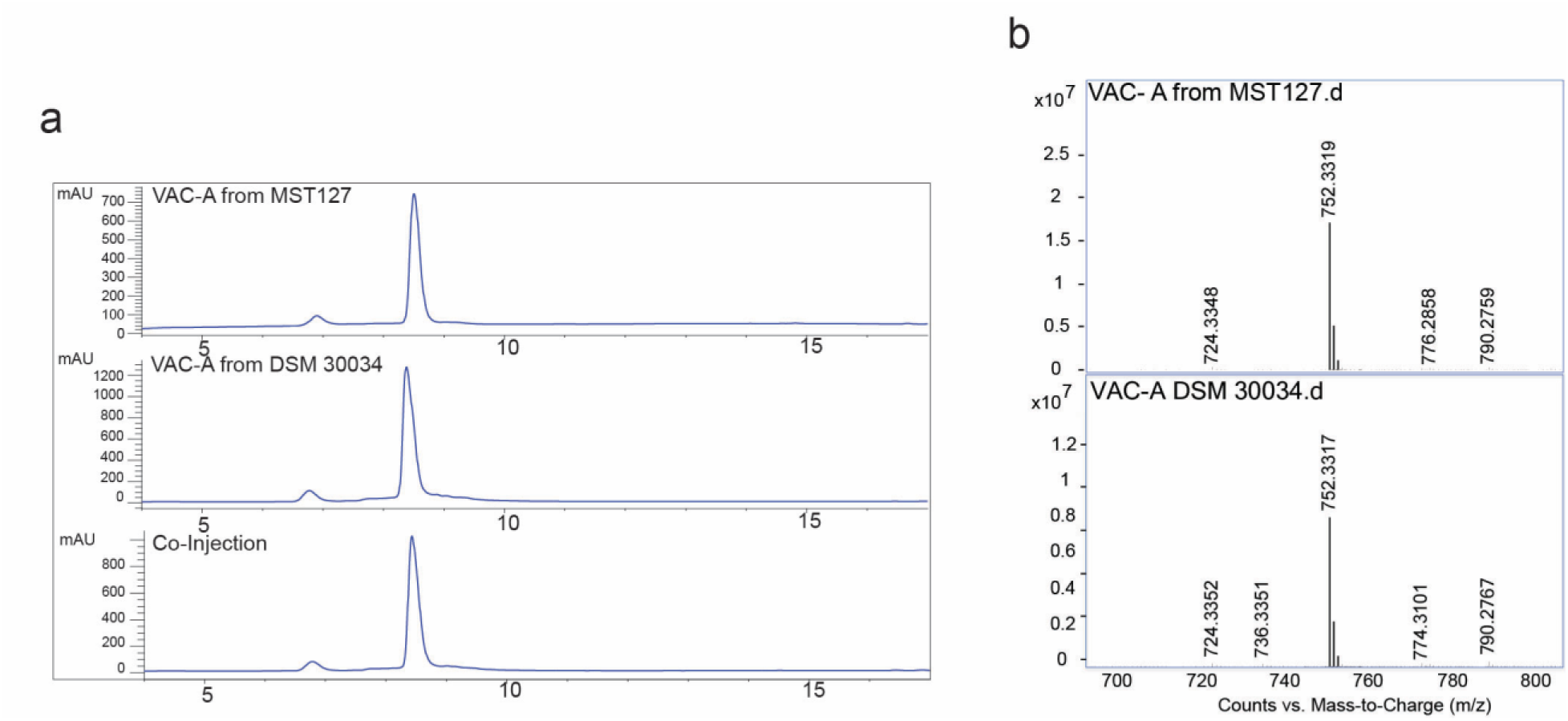
(a) HPLC chromatograms of purified Vac A from *Variovorax* strains MST 127 and DSM 30034, along with their co-injection, demonstrate that they are identical molecules. (b) Mass spectra showing the [M+H]^+^ ion at m/z 752.33 for VAC A from both strains.

**Figure S3.**
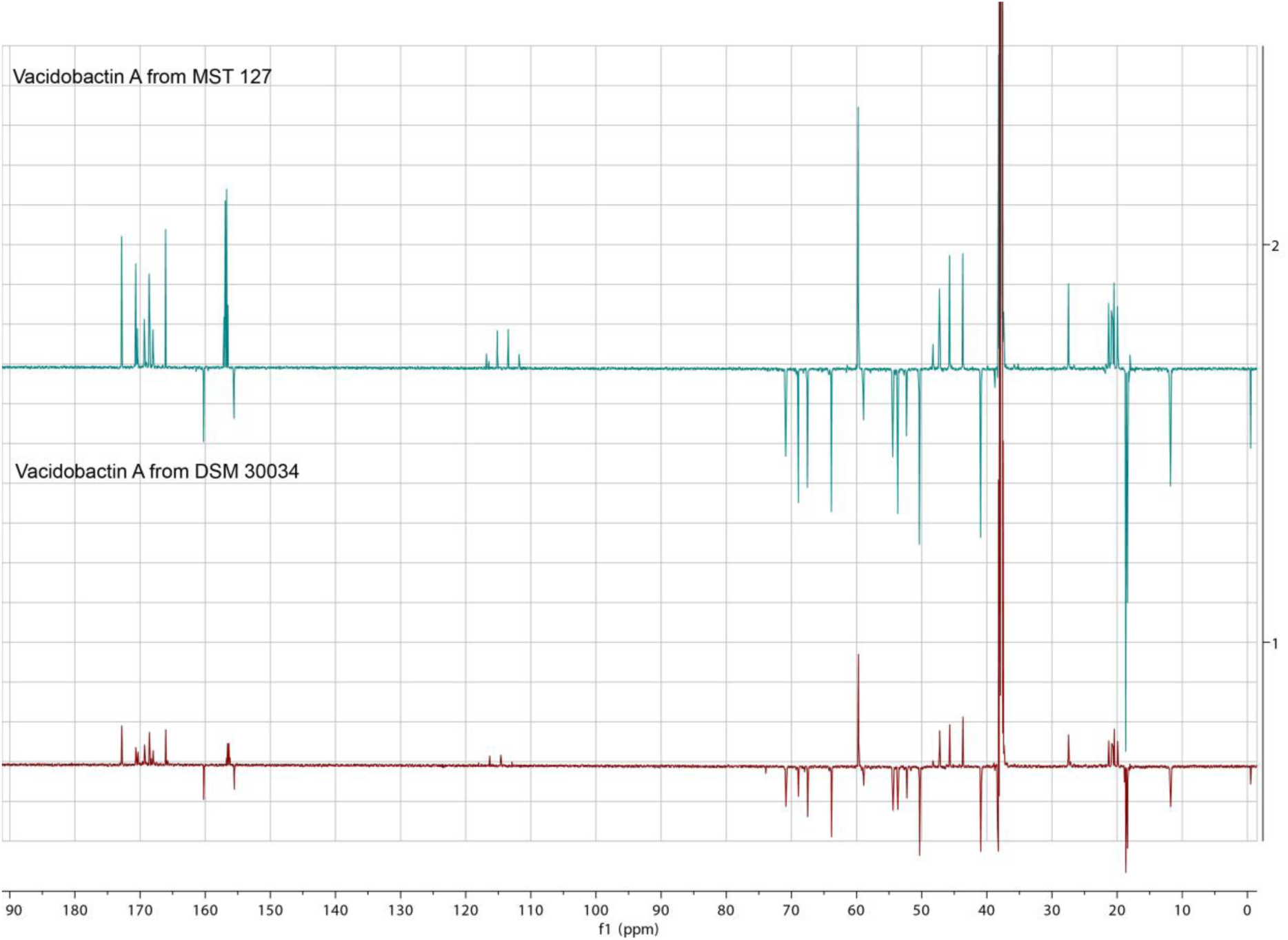
^1^H NMR spectrum of VacA comparison from MST 127 and DSM 30034

**Figure S4.**
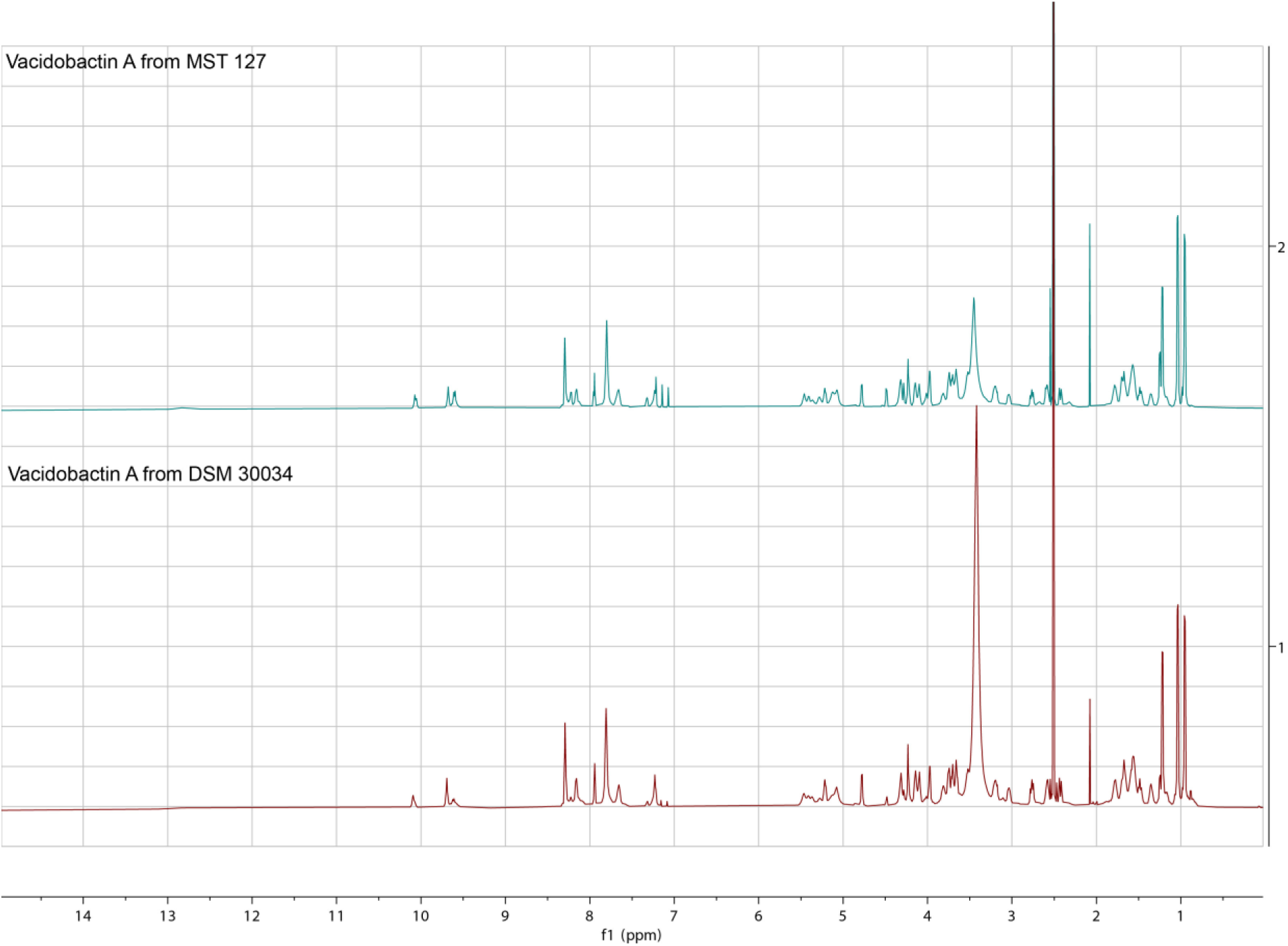
^1^H NMR of Dept-Q spectrum of VacA comparison 1 from MST 127 and DSM 30034

**Figure S5.**
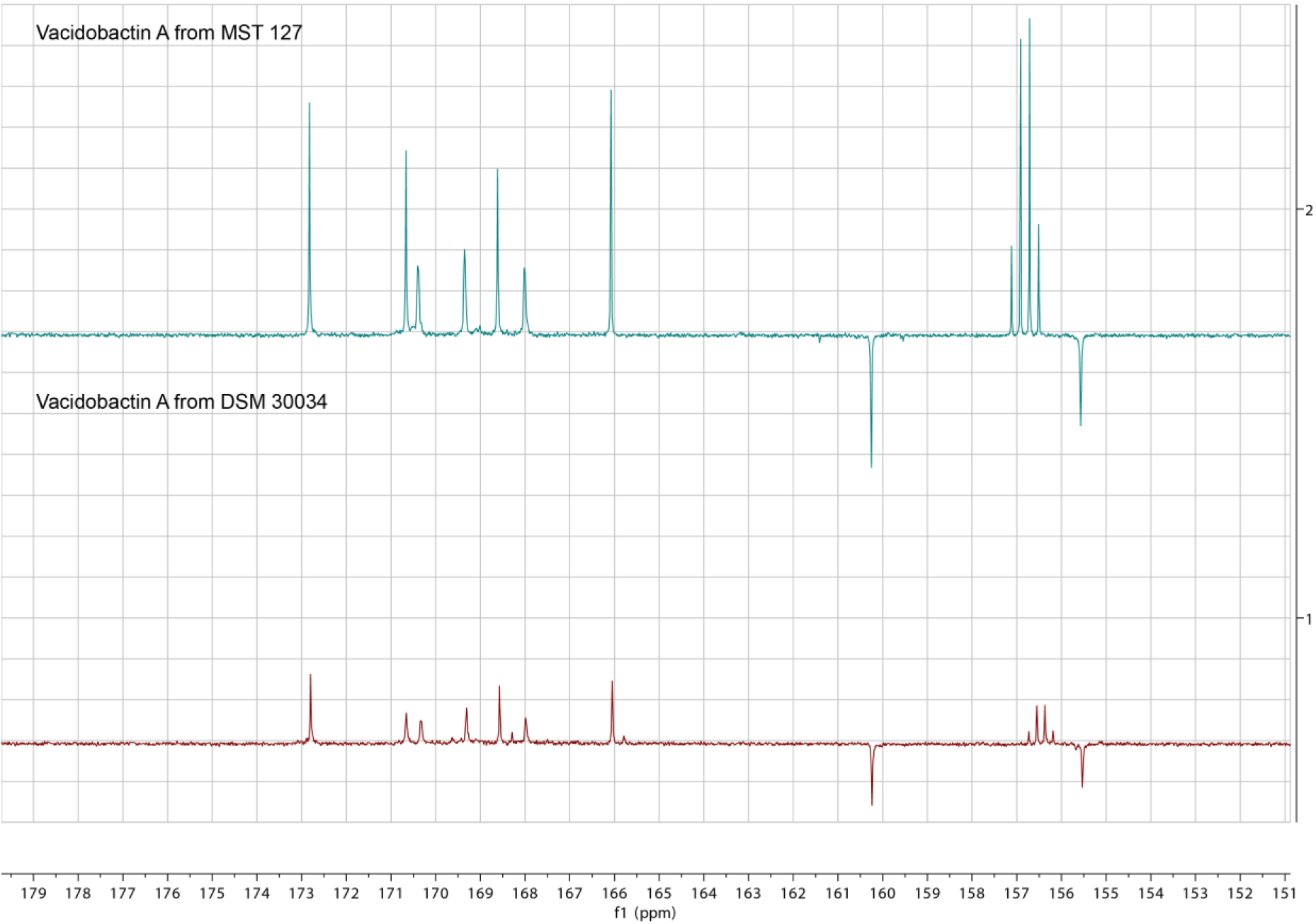
^1^H NMR of Dept-Q spectrum of VacA comparison 2 from MST 127 and DSM 30034

**Figure S6.**
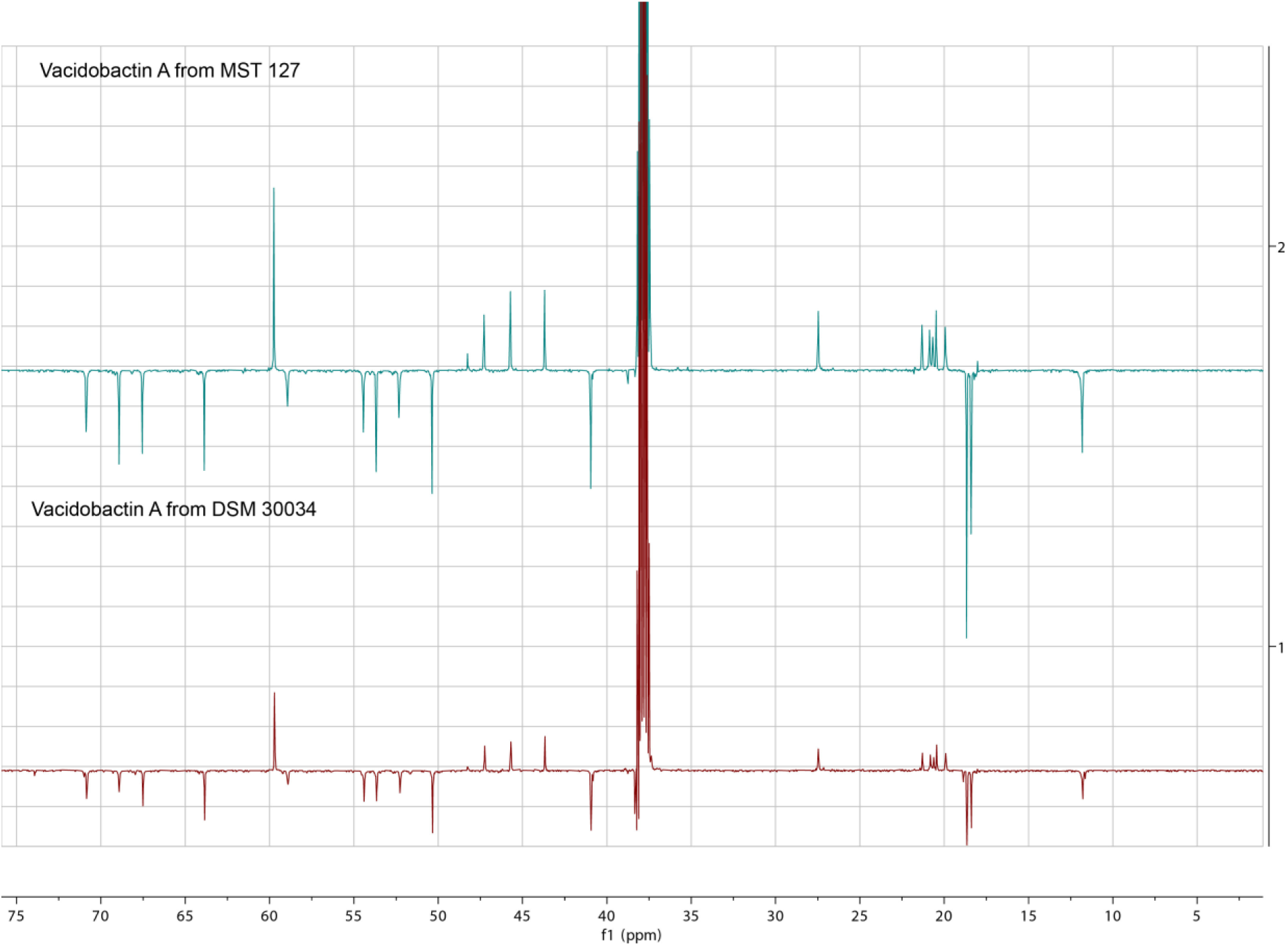
^1^H NMR of Dept-Q spectrum of VacA comparison 3 from MST 127 and DSM 30034

**Figure S7.**
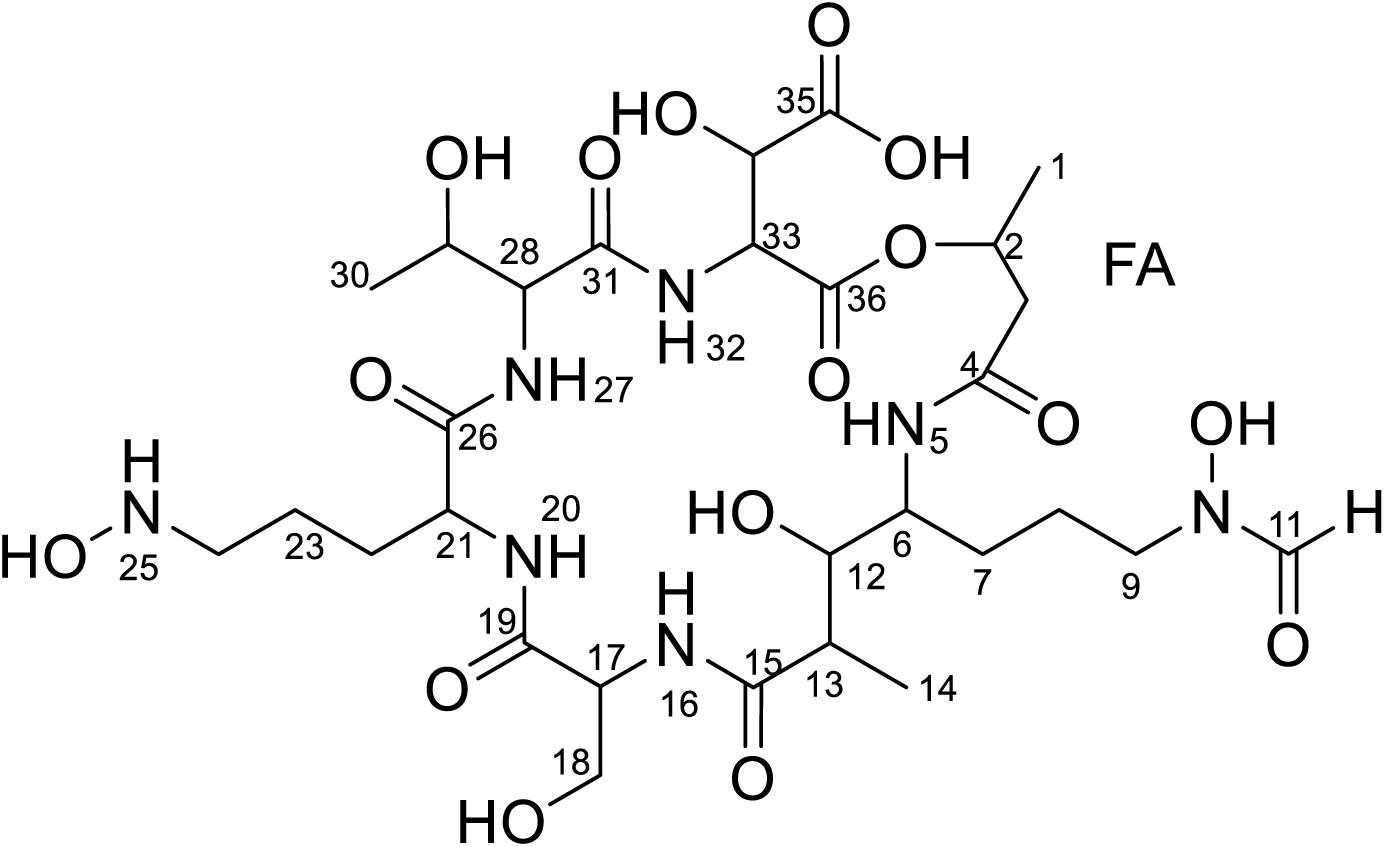
Structural assignments of Vacidobactin A from MST127

**Figure S8.**
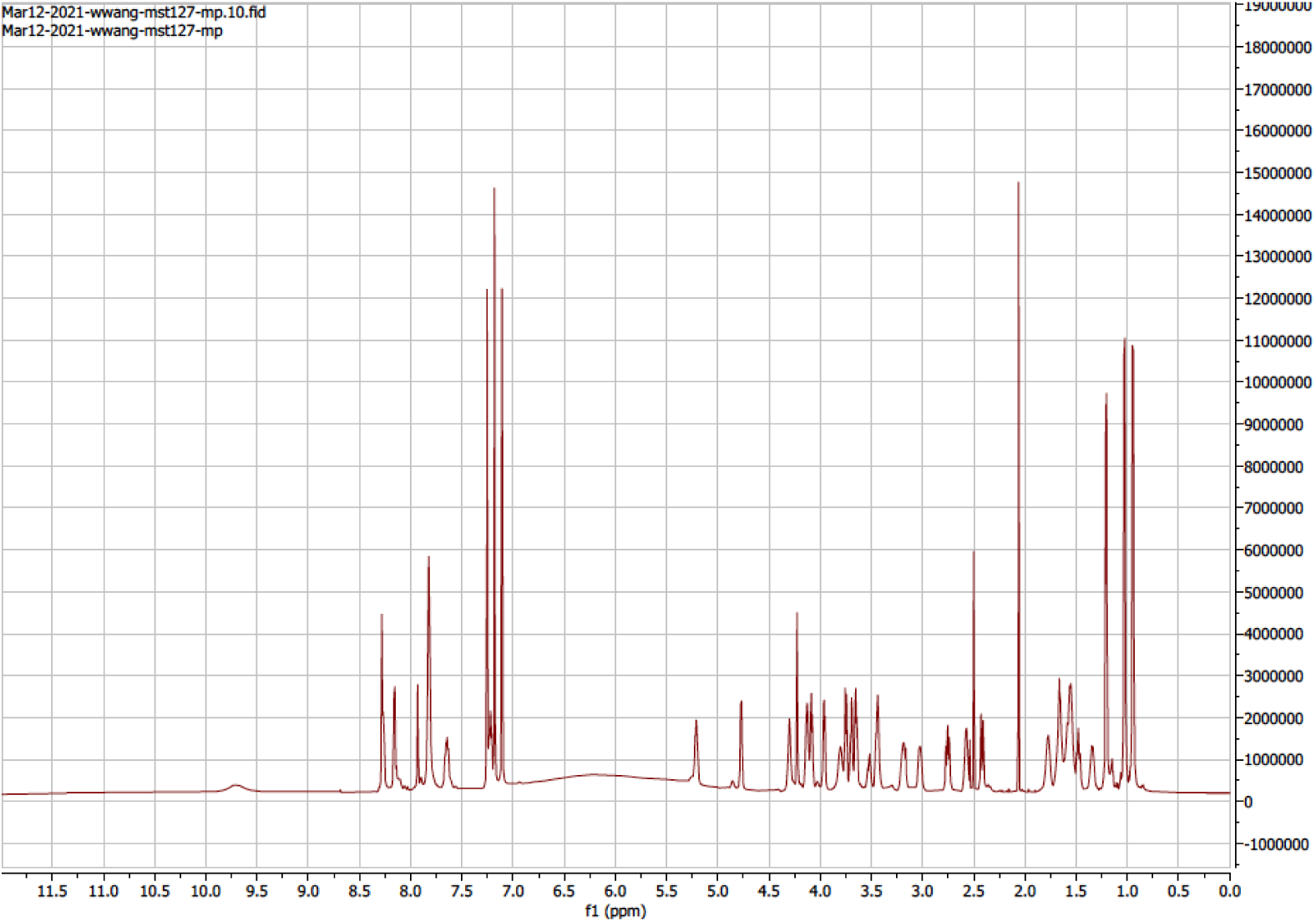
^1^H NMR spectrum of VacA from MST127

**Figure S9.**
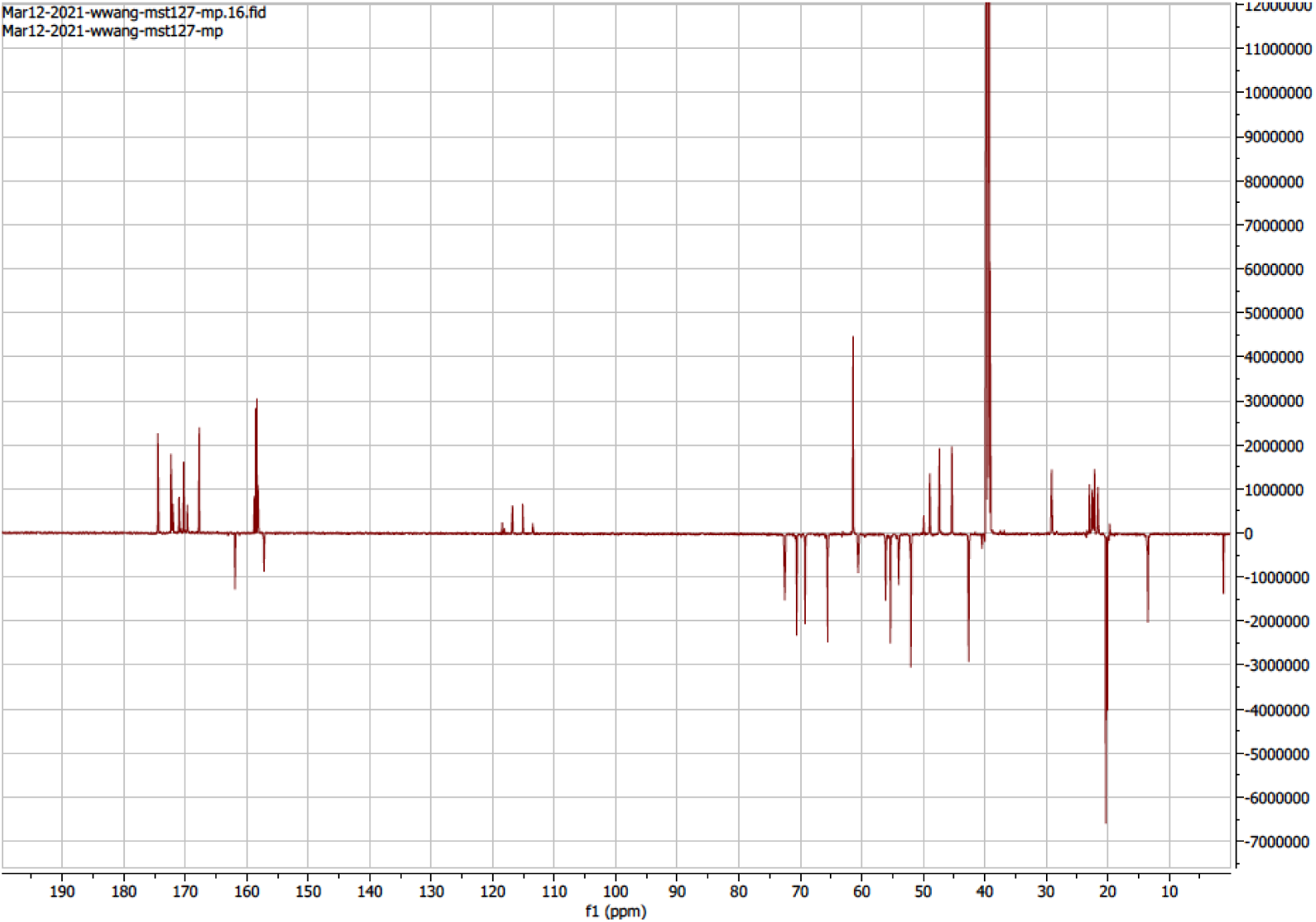
^13^C DEPTQ NMR spectrum of VacA from MST127

**Figure S10.**
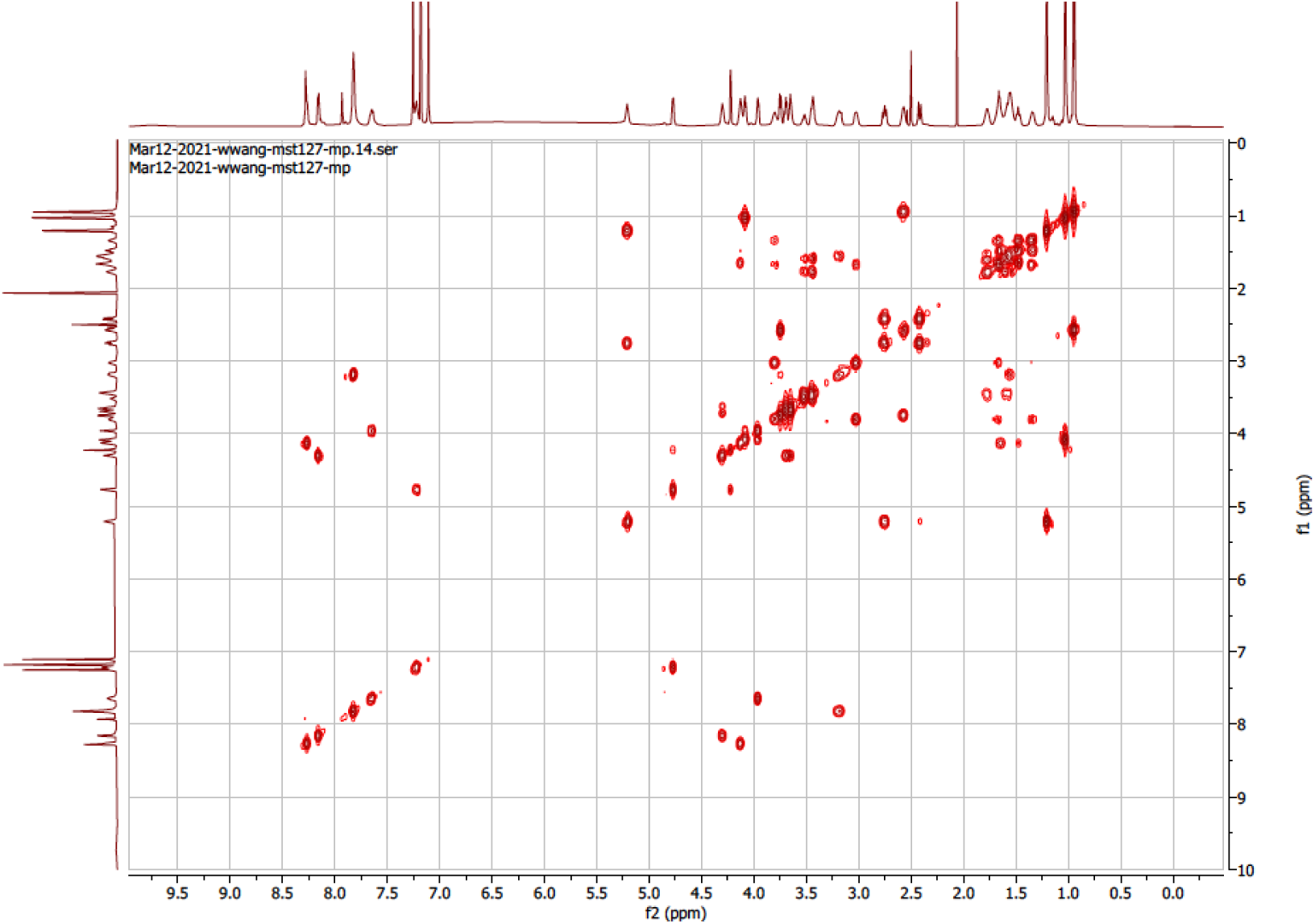
^1^H-^1^H COESY NMR spectrum of VacA from MST127

**Figure S11.**
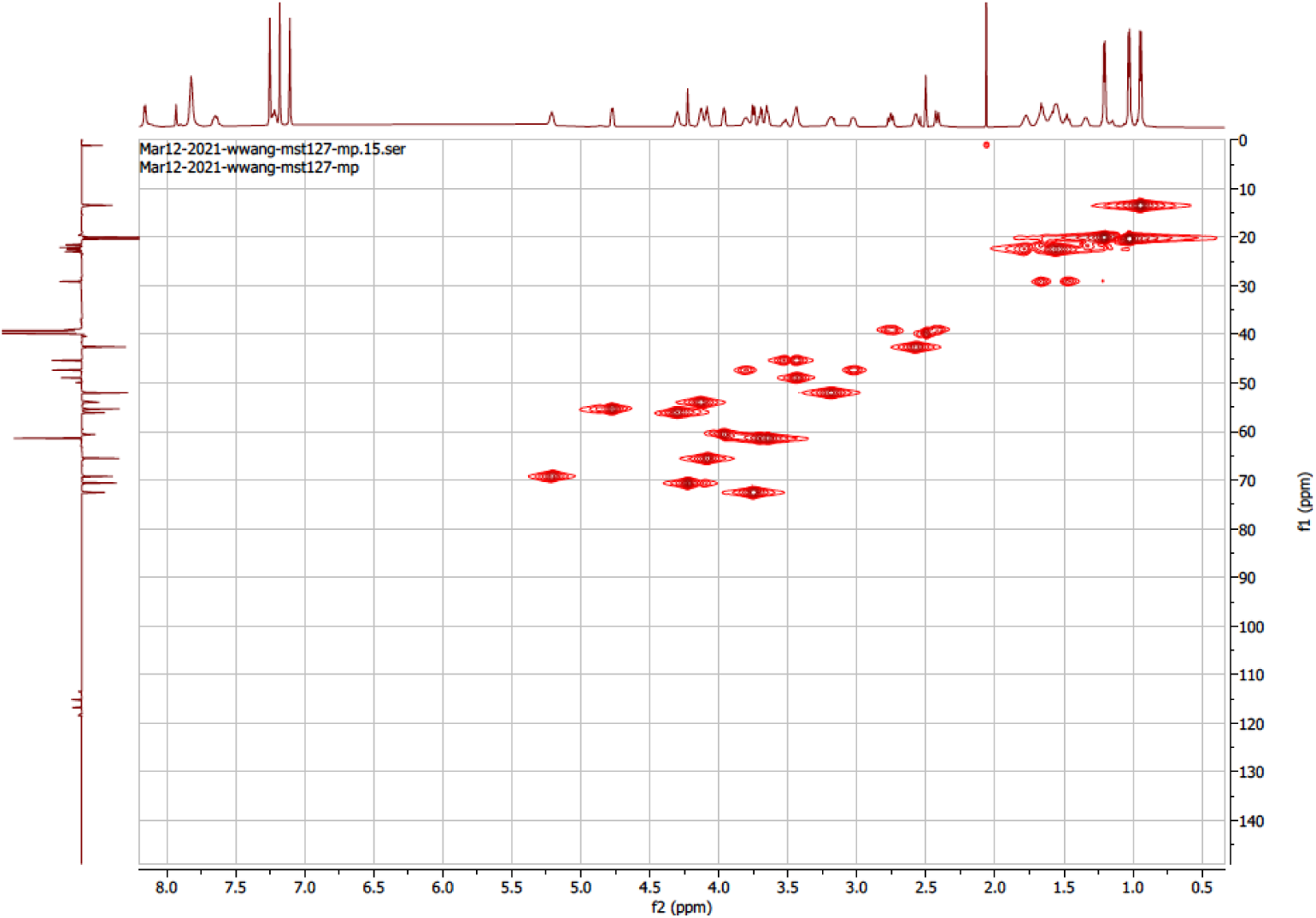
^1^H-^13^C HSQC NMR spectrum of VacA from MST127

**Figure S12.**
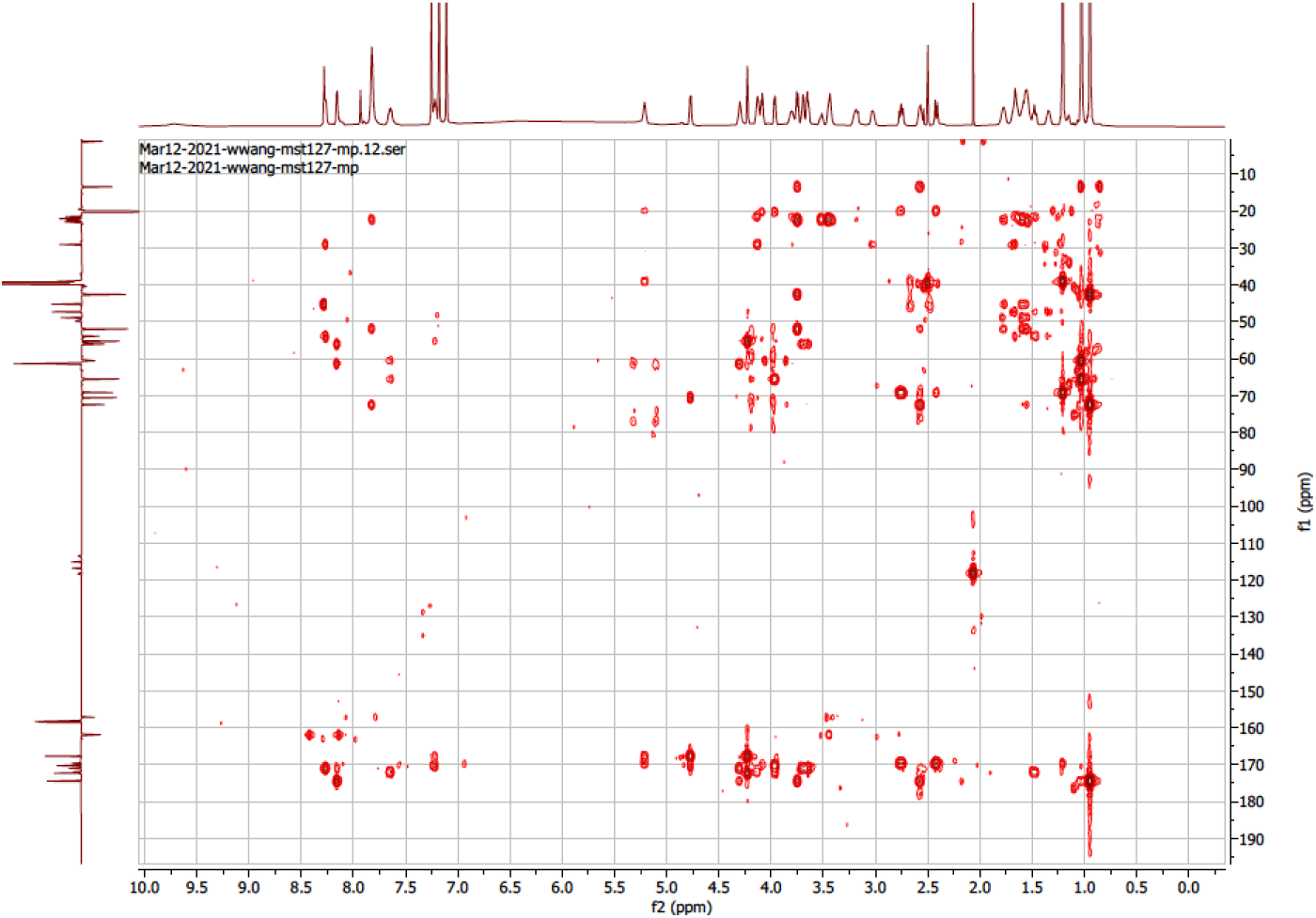
^1^H-^13^C HMBC NMR spectrum of VacA from MST127

**Figure S13.**
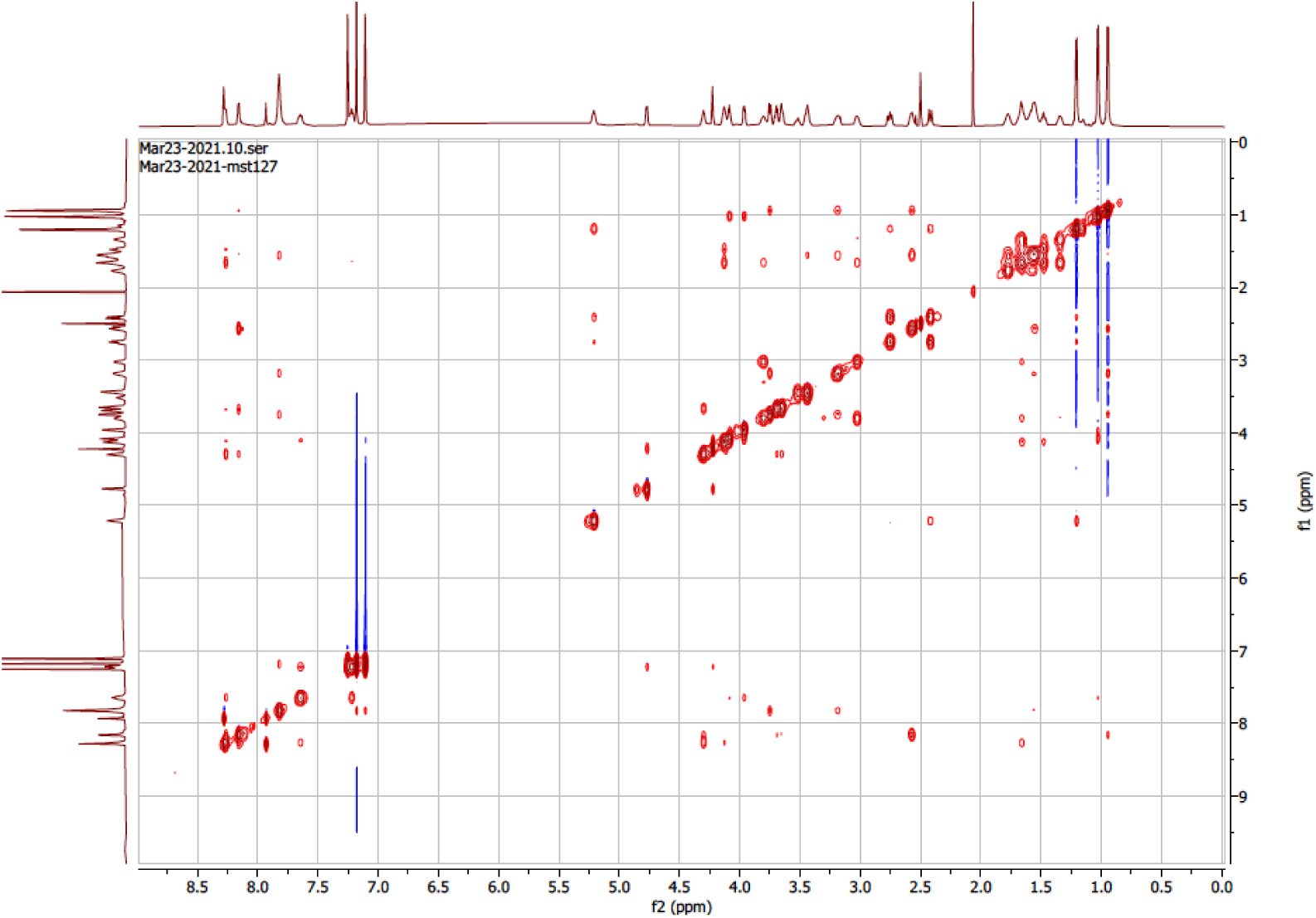
NOESY-NMR spectrum of VacA from MST127

**Figure S14.**
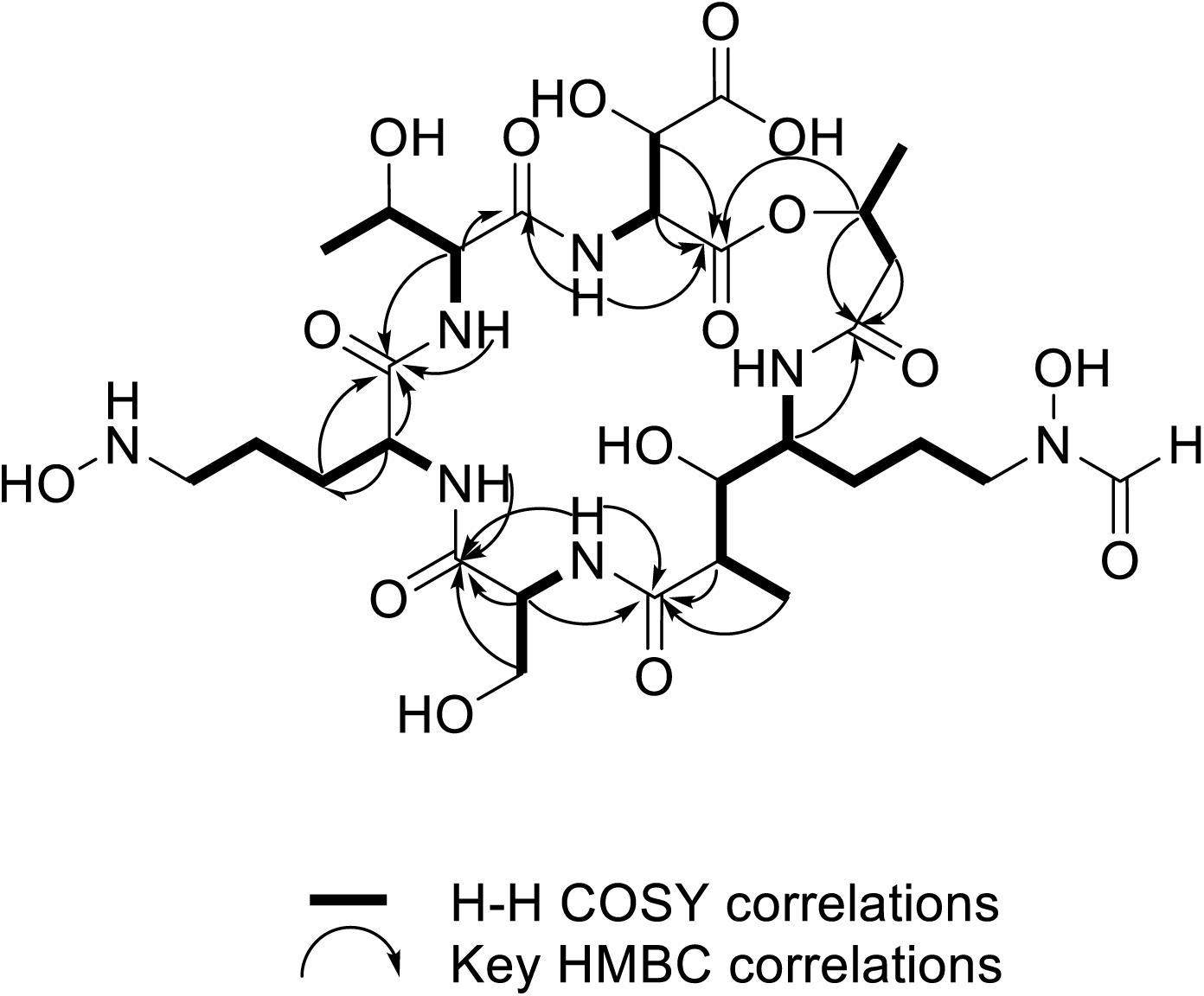
Key COSY- and HMBC-NMR correlations used for the structural elucidation of VacA from MST127.

**Figure S15.**
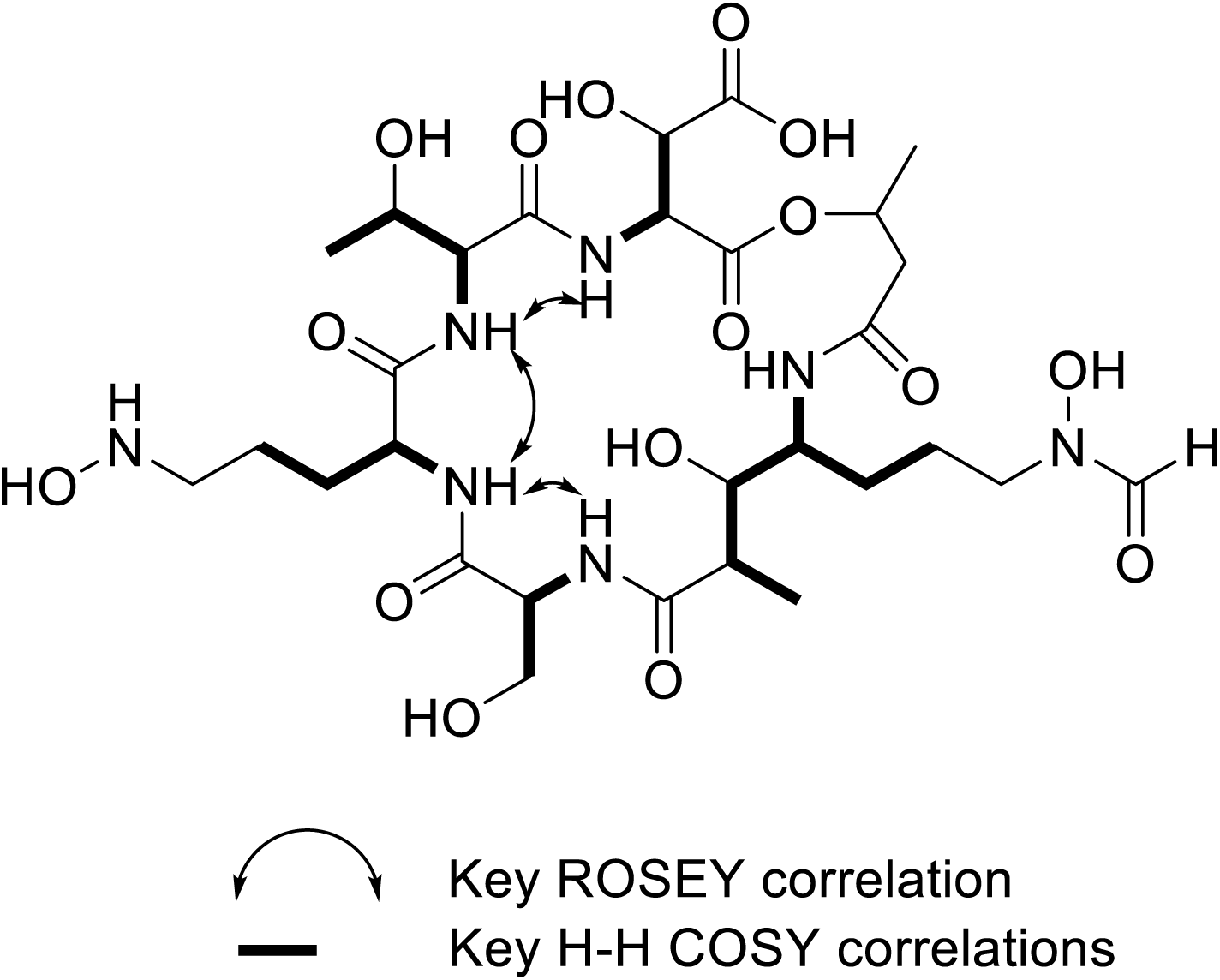
Key ROSEY- and COSY-NMR correlations used for the structural elucidation of VacA from MST127.

**Figure S16.**
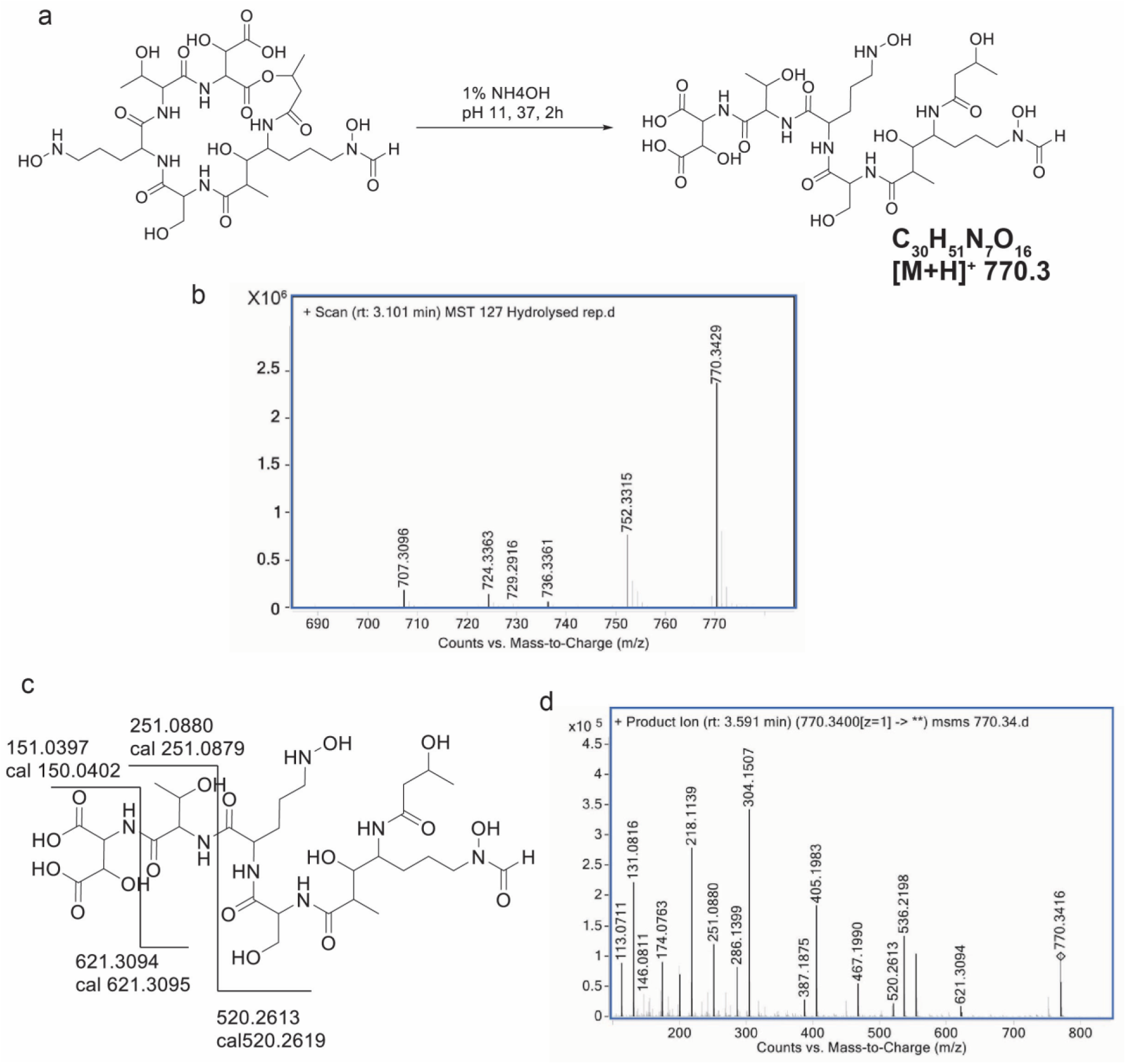
(a) Hydrolysis of VacA leading to ring opening. (b) Confirmation of the hydrolyzed product by mass shift: addition of 18 Da to the parent ion ([M+H] ^+^ 752.33) yields [M+H]^+^ 770.34. (c) MS/MS fragmentation pattern and structural interpretation of intact VacA. (d) MS/MS fragmentation spectrum of hydrolyzed Vac-A, confirming the proposed fragmentation pattern.

**Figure S17.**
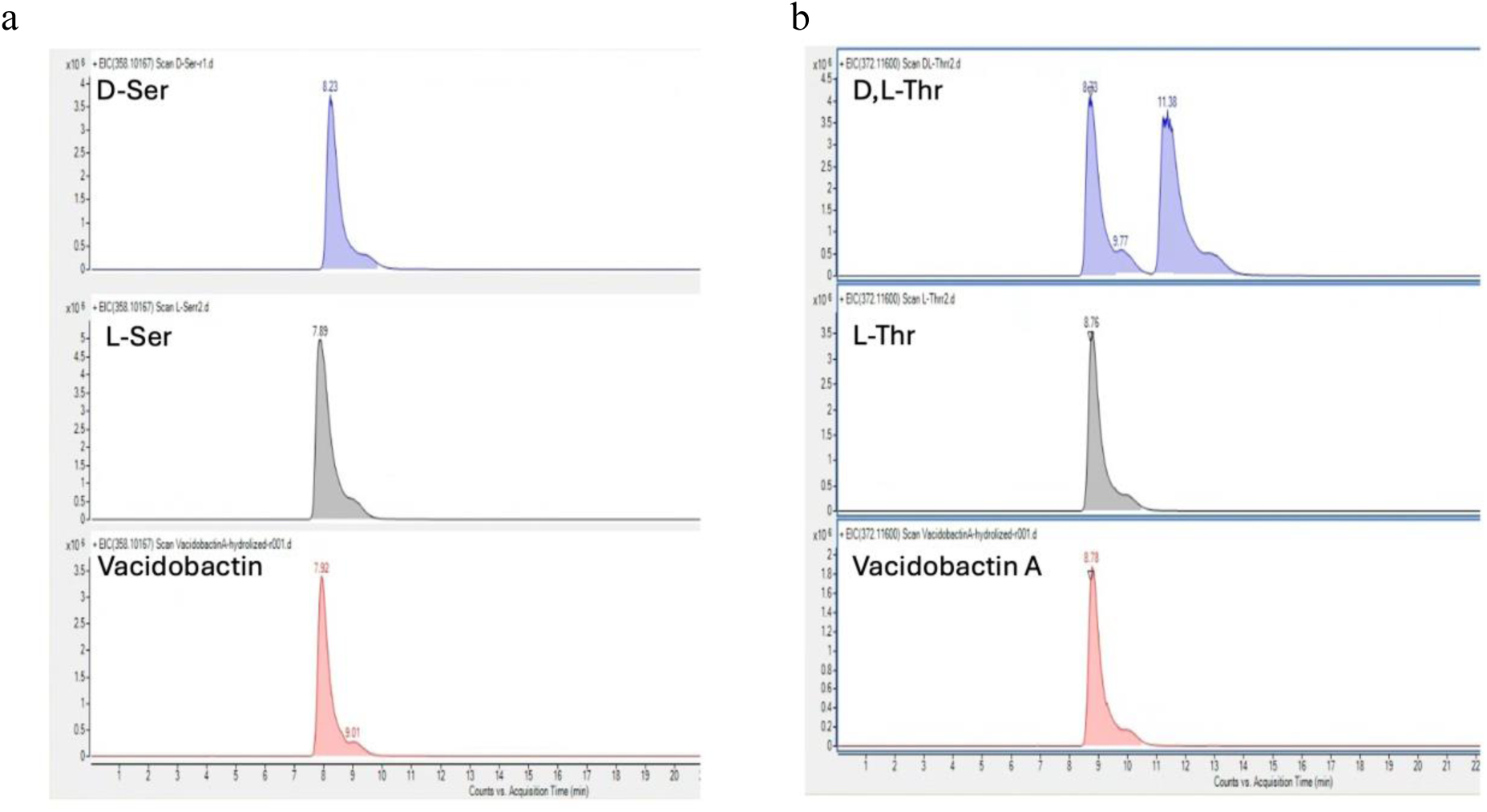
a) Extracted ion Chromatograms (EIC) at 358.10167, corresponding to derivatized serine; b). EIC at 372.11690, corresponding to the modified threonine. The other 3 non-proteinogenic amino acids (4-amino-3-hydroxy-7-(N-hydroxyformamido)-2-methylheptanoic acid, 2-amino-5-(hydroxyamino)pentanoic acid, and β-hydroxyaspartic acid) could not be detected using this approach.

**Figure S18.**
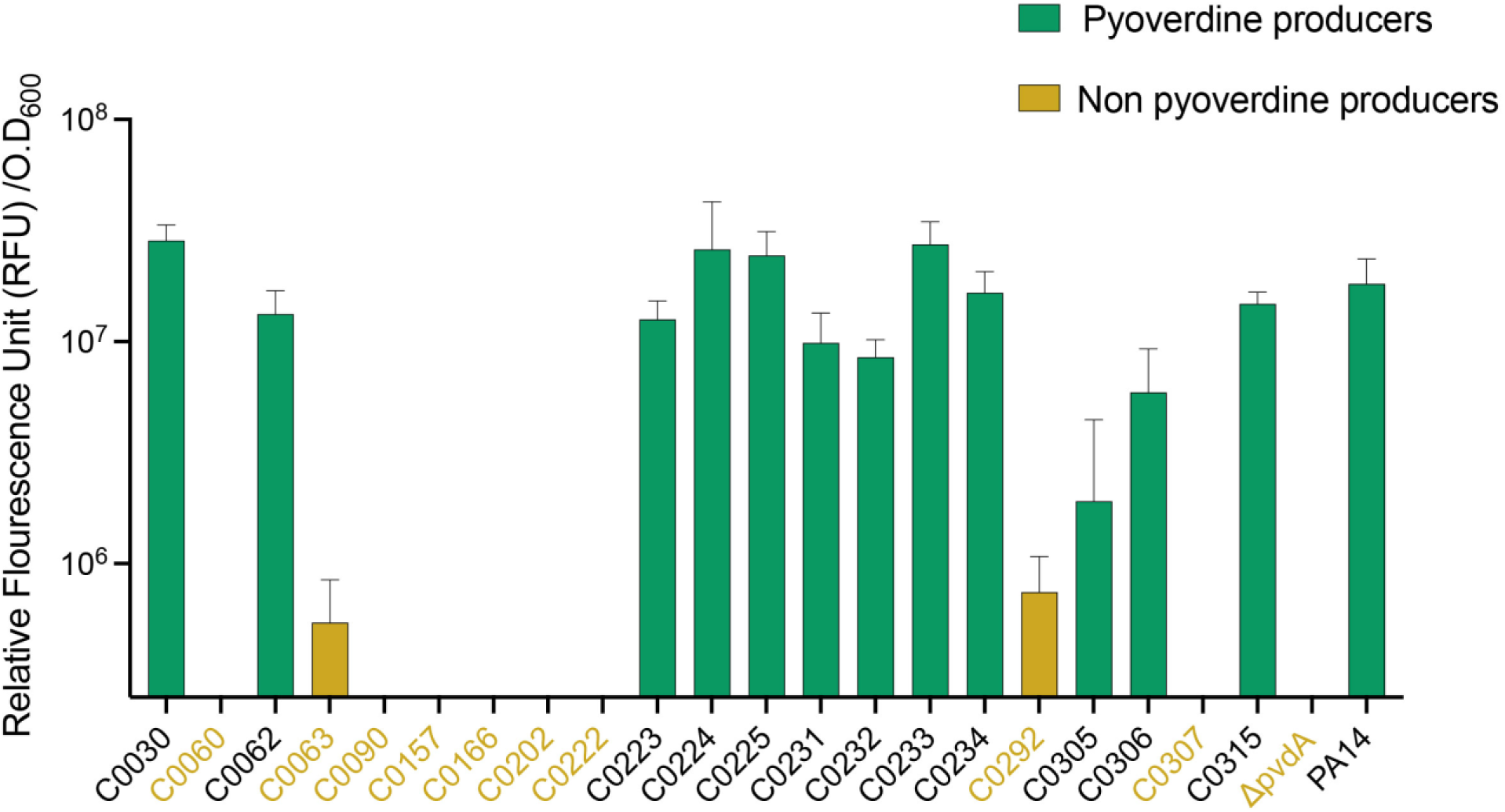
Pyoverdine production among *P. aeruginosa* strains from the Wright clinical collection. Pyoverdine producers are indicated in green, and non-producers in yellow. The assay was performed in triplicate with three independent biological replicates. Optical density at 600 nm (OD₆₀₀) was consistent across all isolates, indicating that differences in fluorescence reflect true variation in siderophore production rather than growth differences.

**Figure S19.**
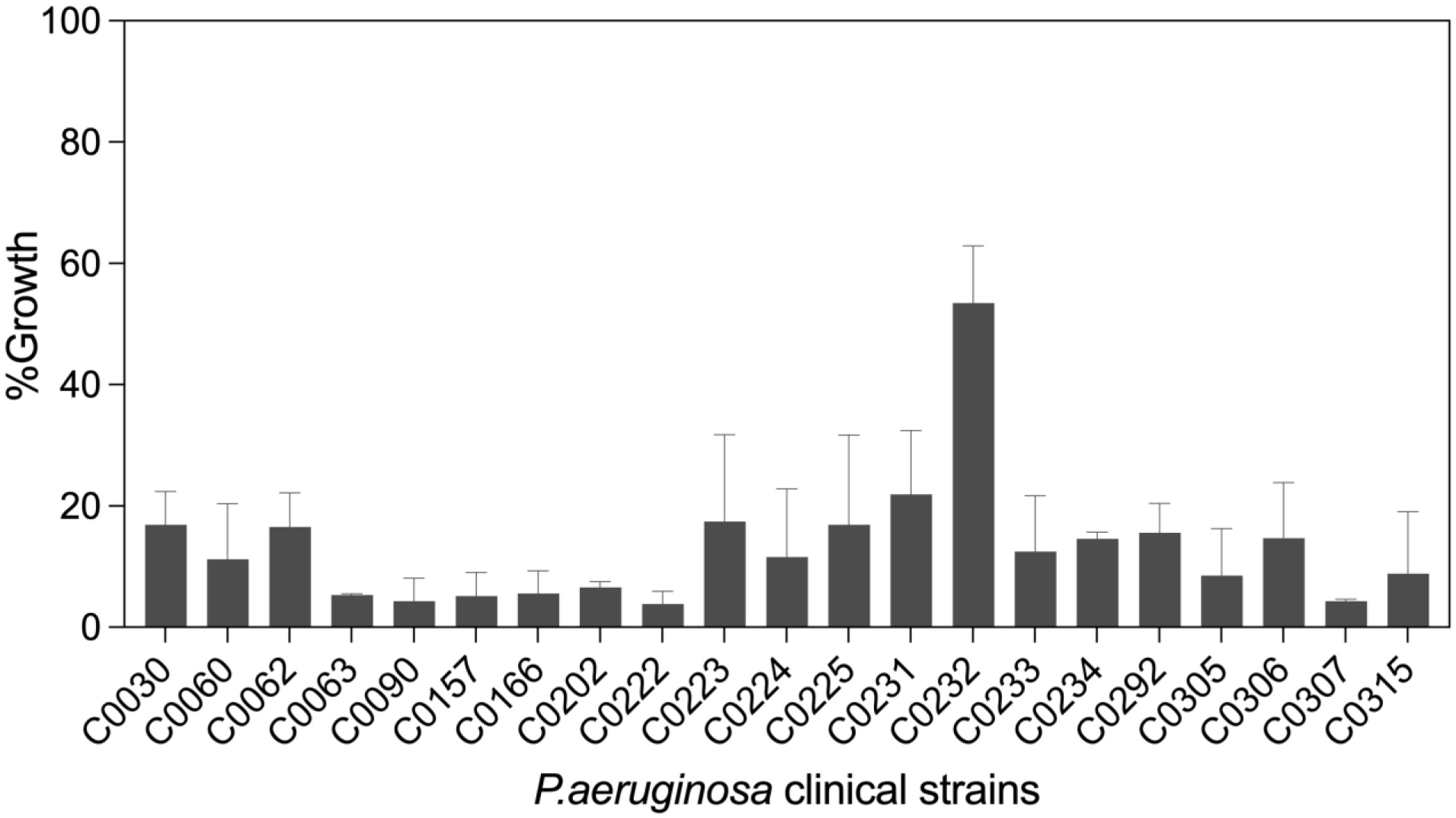
Bar graph representing the combination assay results against various clinical isolates of *P. aeruginosa*. The combination of VacA (8 µg/mL) and thiostrepton (0.5 µg/mL) was used. Percent growth (y-axis) is shown for each clinical strain (x-axis) in the presence of the combined treatment. The dashed line indicates that growth was reduced to below 20% in many strains, demonstrating synergy. Error bars represent the standard deviation from triplicate experiments. The error bars suggest variability in growth measurements among replicates, particularly for isolates with higher percent growth.

## SUPPLEMENTARY TABLES

**Table S1.**
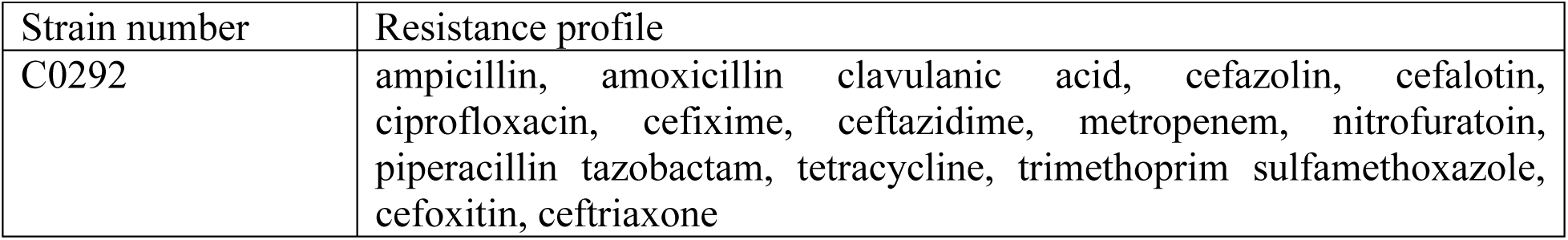
Antibiogram of WCC strain C0292 *P. aeruginosa* used for screening for antimicrobial activity.

**Table S2.**
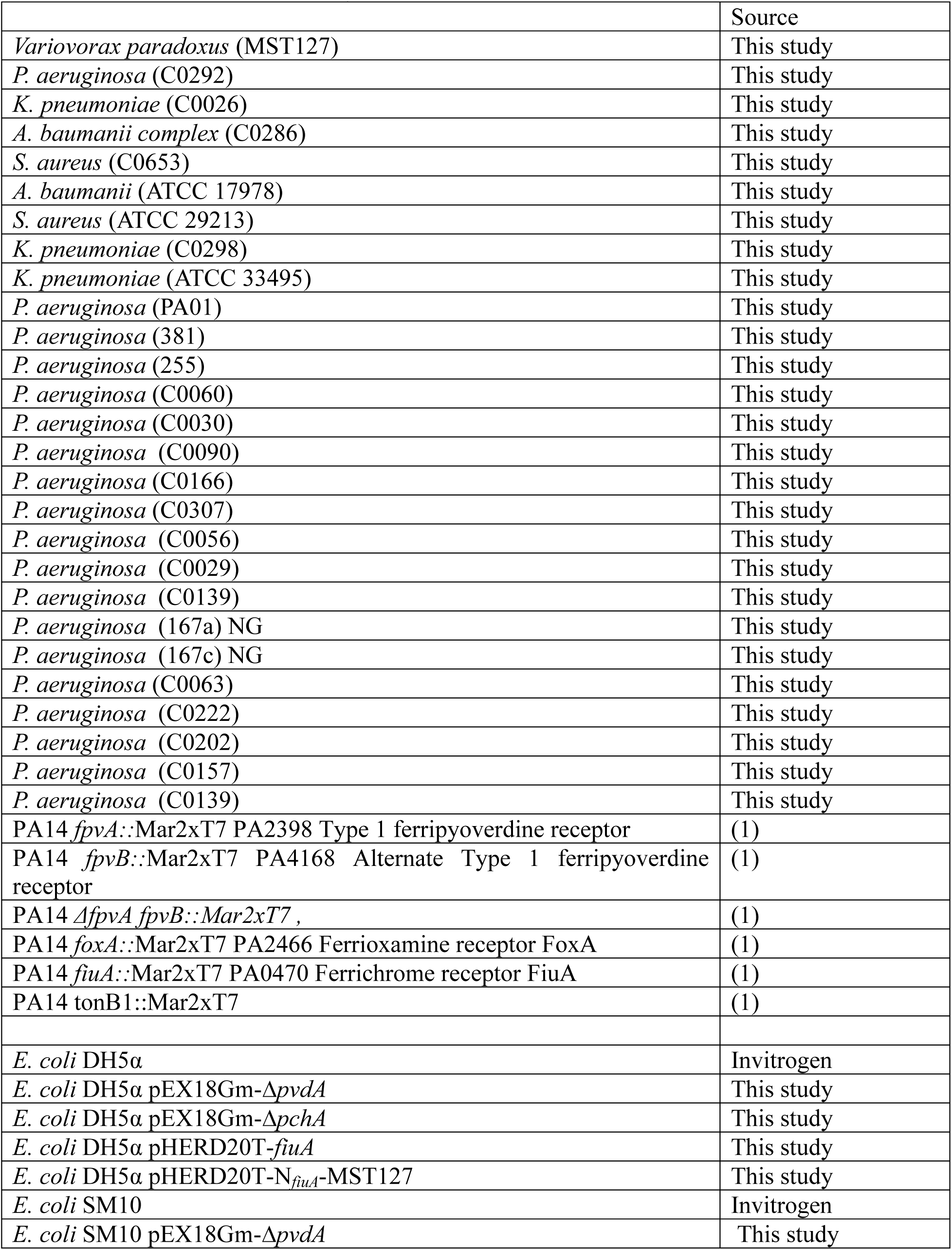
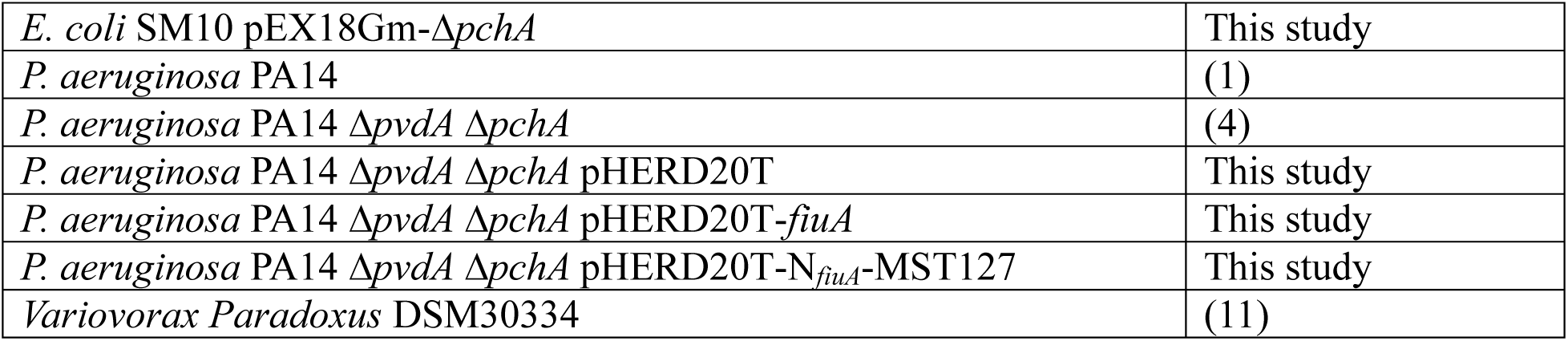
Strains used in this study.

**Table S3.** Plasmids used in this study.

|  | Source |
| --- | --- |
| pEX18Gm | (2) |
| pEX18Gm-$\Delta pvdA$ | This study |
| pEX18Gm-$\Delta pchA$ | This study |
| pHERD20T | (3) |
| pHERD20T-*f<sub>iuA</sub>* | This study |
| pHERD20T-N<sub>*f<sub>iuA</sub>*</sub>-MST127 | This study |

**Table S4.**
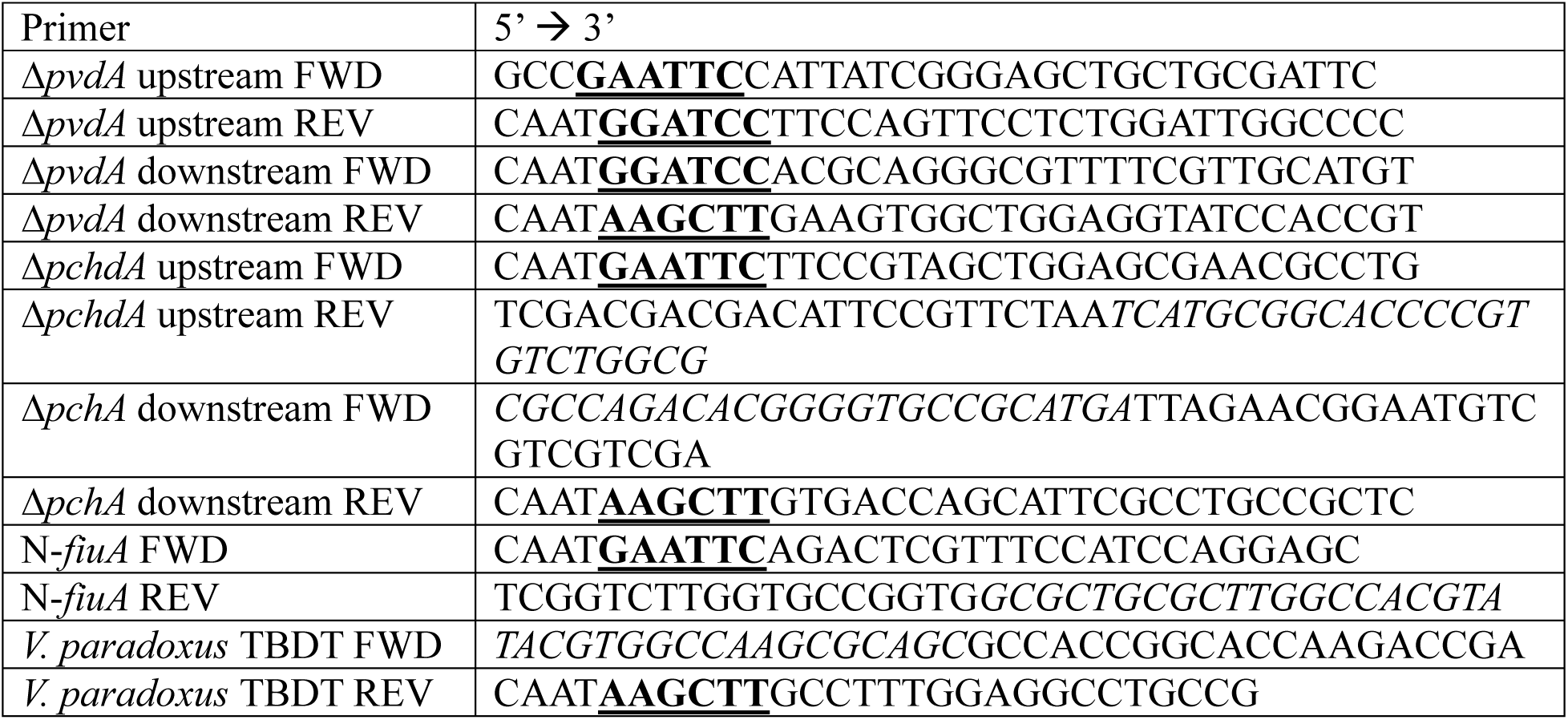
Primers used in this study. Restriction enzyme sites are bolded and underlined. Overlapping regions are italicized.

**Table S5.**
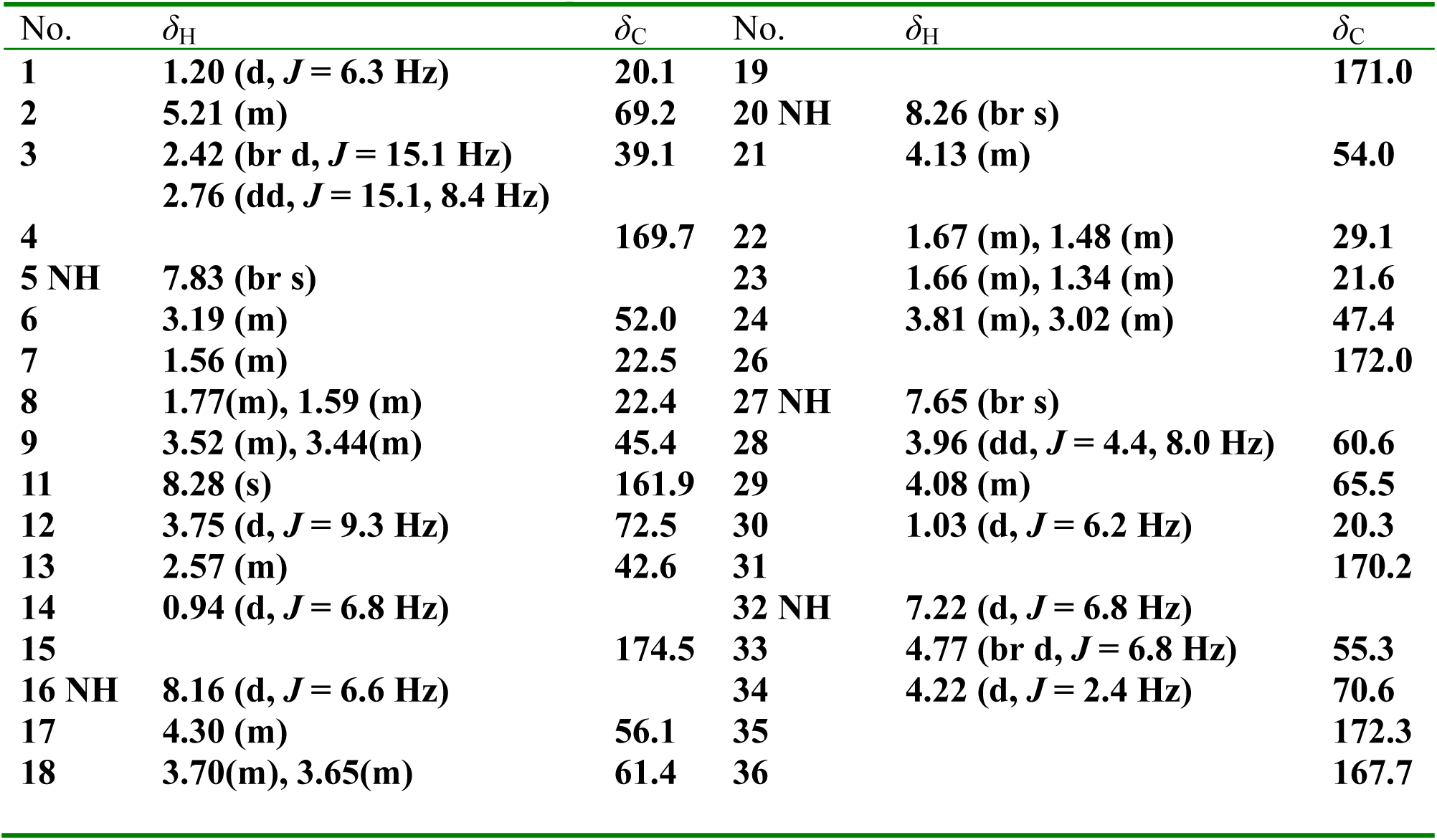
H and C NMR Data of compound Vac A from MST127.

**Table S6.** The comparison of between the genes and their putative functions from BGCs.

| **Vacidobactin gene cluster analysis**<br>**(*Variovorax paradoxus* S110 (DSM No. 30034))** |  |  |  | **Vacidobactin gene cluster analysis**<br>**(*Variovorax paradoxus* MST 127)** |  |  |  |  |
| --- | --- | --- | --- | --- | --- | --- | --- | --- |
| **Locus** | **Predicted Function** | **Strand** | **Amino Acids** | **Locus** | **Location** | **Predicted Function** | **Strand** | **Amino acids** |
| Vapar_3752 | RNA polymerase, sigma-24 subunit, ECF subfamily | - | 181 | ctg8_230 | 248,867 - 249,412 | Probable RNA polymerase sigma factor FecI | - | 181 |
| Vapar_3751 | anti-FecI sigma factor | - | 67 | ctg8_229 | 248,567 - 248,770 | no-hits | - | 67 |
| Vapar_3750 | MbtH domain protein | - | 85 | ctg8_228 | 248,209 - 248,466 | mbtH-like protein | - | 85 |
| Vapar_3749 | Thioesterase | - | 246 | ctg8_227 | 247,466 - 248,203 | thioesterase | - | 245 |
| Vapar_3748 | Phosphopantetheinyl transferase | - | 229 | ctg8_226 | 246,799 - 247,488 | 4'-phosphopantetheinyl transferase | - | 229 |
| Vapar_3747 | TauD dioxygenase | - | 330 | ctg8_225 | 245,810 - 246,802 | Dioxygenase TauD/TfdA | - | 330 |
| Vapar_3746 | NRPS | - | 1776 | ctg8_224 | 240,474 - 245,789 | NRPS | - | 1771 |
| Vapar_3745 | PKS | - | 1520 | ctg8_223 | 235,912 - 240,474 | PKS | - | 1520 |
| Vapar_3744 | NRPS | - | 1110 | ctg8_222 | 232,574 - 235,915 | NRPS | - | 1113 |
| Vapar_3743 | NRPS | - | 2626 | ctg8_221 | 224,697 - 232,574 | NRPS | - | 2625 |
| Vapar_3742 | NRPS | - | 1358 | ctg8_220 | 220,627 - 224,700 | NRPS | - | 1357 |
| Vapar_3741 | TonB siderophore receptor | + | 723 | ctg8_219 | 218,377 - 220,545 | TonB-dependent Receptor | + | 722 |
| Vapar_3740 | L-lysine 6-monooxygenase | + | 439 | ctg8_218 | 216,862 - 218,214 | lysine/ornithine N-monooxygenase | + | 450 |
| Vapar_3739 | N-acetyltransferase | + | 344 | ctg8_217 | 215,770 - 216,810 | Acetyltransferase (GNAT) | + | 346 |
| Vapar_3738 | N5-hydroxyornithine formyltransferase | + | 281 | ctg8_216 | 214,928 - 215,773 | hydroxymethyl-, formyl- and related transferase activity | + | 302 |
| Vapar_3737 | Ferric iron reductase | + | 281 | ctg8_215 | 214,047 - 214,826 | Ferric iron reductase<br>FhuF-like<br>transporter | + | 259 |
| Vapar_3736 | Cyclic peptide<br>transporter | - | 563 | ctg8_214 | 212,151 - 213,842 | ABC_membran<br>e,<br>ABC_transport<br>er | - | 563 |

